# Gastruloid-derived Primordial Germ Cell-like Cells (Gld-PGCLCs) develop dynamically within integrated tissues

**DOI:** 10.1101/2023.06.15.545059

**Authors:** Christopher B. Cooke, Christopher Barrington, Peter Baillie-Benson, Jennifer Nichols, Naomi Moris

## Abstract

Primordial Germ Cells (PGCs) are the early embryonic precursors of gametes - sperm and egg cells. PGC-like cells (PGCLCs) can currently be derived *in vitro* from pluripotent cells exposed to signalling cocktails and aggregated into large embryonic bodies, but these do not recapitulate the native embryonic environment during PGC formation. Here we show that mouse gastruloids, a three-dimensional *in vitro* model of gastrulation, contain a population of Gastruloid-derived PGC-like cells (Gld-PGCLCs) that resemble early PGCs *in vivo*. Importantly, the conserved organisation of mouse gastruloids leads to coordinated spatial and temporal localisation of Gld-PGCLCs relative to surrounding somatic cells, even in the absence of specific exogenous PGC-specific signalling or extraembryonic tissues. In gastruloids, self-organised interactions between cells and tissues, including the endodermal epithelium, enables the specification and subsequent maturation of a pool of Gld-PGCLCs. As such, mouse gastruloids represent a new source of PGCLCs *in vitro* and, due to their inherent co-development, serve as a novel model to study the dynamics of PGC development within integrated tissue environments.

---

The specification of mouse Primordial Germ Cells (PGCs) occurs at the gastrulation-stage epiblast at about embryonic day (E)7.25, where competent cells begin to co-express *Stella* and *Blimp1* and become lineage-restricted to a germ cell fate^1-4^ by repression of somatic genes and the activation of the PGC-specific program^5,6^. This specification occurs at the proximal posterior of the epiblast, and is thought to be dependent on signals from the Extraembryonic Ectoderm (ExE) and Visceral Endoderm (VE), including BMP^2^ and Wnt signalling^7^, since embryos mutant for *Bmp4* or one of its receptors, *ALK2,* have reduced numbers of PGCs^8,9^. After specification, PGCs are incorporated into the developing hindgut, and move anteriorly through this tissue before then migrating through the dorsal mesentery towards the genital ridge^10,11^, the precursors of the gonads. Here, the germ cells colonise the prospective gonadal niche in the form of small cell clusters^10^, and continue to mature in terms of their transcriptional and, particularly, their epigenetic signature. At approximately E12.5^12,13^ sexual determination occurs, and initiates further sex-specific maturation that ultimately generates spermatozoa in males and oocytes in females. Their time-course is therefore highly dynamic, and occurs through close association with several different tissues and cell types of the developing embryo^14^.

Currently, pluripotent stem cell-based PGC-like cell (PGCLC) *in vitro* models^15,16^, have been used to explore the regulatory mechanisms of early specification and maturation (for example, ^17,18^) and even to generate mature germ cells through gametogenesis^15,19-24^. These models are typically derived from Epiblast-like cells (EpiLCs) which are subsequently arranged as embryoid bodies, and they build on earlier work that observed spontaneous PGCLC differentiation in EBs^25-27^, but with the addition of PGC-specific factors to strongly bias towards a PGCLC fate. Yet, despite being an efficient protocol, these EB-derived PGCLCs are formed within largely disorganised aggregates of cells that lack the spatially-organised, supportive neighbouring cell types found in the embryo, and have limited epigenetic remodelling towards mature germ cells^28,29^. In addition, further maturation of PGCLCs beyond the gonadal colonisation stage *in vitro* currently requires complete dissociation of EBs and reaggregation with gonadal cell populations^19-21,30^, which necessarily results in loss of any endogenous spatial colocalization or organisation and precludes any study of the gradual developmental dynamics of PGCLCs during this maturation time window. Therefore, while embryoid body-based methods provide a readily available source of *in vitro* PGCLCs, these methods are unable to reveal the complexities of PGC specification or their interaction with the rest of the embryonic body plan in a developmentally faithful manner.

Recently, mouse gastruloids, three-dimensional mouse embryonic stem (ES) cell-derived aggregates, have been described and characterised to undergo gastrulation-like gene expression progression, multilineage differentiation, axial polarisation, and morphological extension^31,32^. Single-cell analysis showed that these gastruloids include many cell types found in the early mouse embryo, including a population of presumptive PGCLCs^33^. Others have also shown that small populations of *Sox2*+/DPPA3+ cells^34^ and DPPA4+ cells^35^ exist along the anteroposterior length of gastruloid-like structure. Here, we report the further characterisation of these gastruloid-derived PGCLCs (Gld-PGCLCs), including their dynamic spatiotemporal localisation and association within integrated tissue environments. Importantly, we show that Gld-PGCLCs display characteristics that are akin to *in vivo* PGCs, and that they recapitulate features of early PGC migration and maturation, reaching stages equivalent to ∼E14.5, while relying mainly on endogenous inductive signals from within the self-organised gastruloid.

## Results

### Identification of mouse gastruloid-derived PGCLCs

The transcriptional expression of *Blimp1* (also known as *Prdm1*) and *Stella* (*Dppa3*) are both associated with PGCs in the mouse embryo^4,36^. We therefore generated mouse gastruloids using the Blimp1:eGFP (herein, Blimp1-GFP)*^4^* and Blimp1:mVenus Stella:CFP (BVSC)^37^ mouse embryonic stem cell lines, which have previously been used as markers of PGCLC state *in vitro^38^*. Aggregates made from BVSC and Blimp1-GFP cells broke symmetry at approximately 96 hours after aggregation (h), leading to elongated structures with polarised expression of the mesodermal marker *Brachyury* (T-BRA) and CDX2 at 120h (Figure 1A-B, Supplementary Figure 1A) comparable to gastruloids generated from E14tg2A cells^31,39^ routinely cultured in 2iLIF (see Methods)^40,41^.

**Figure 1:**
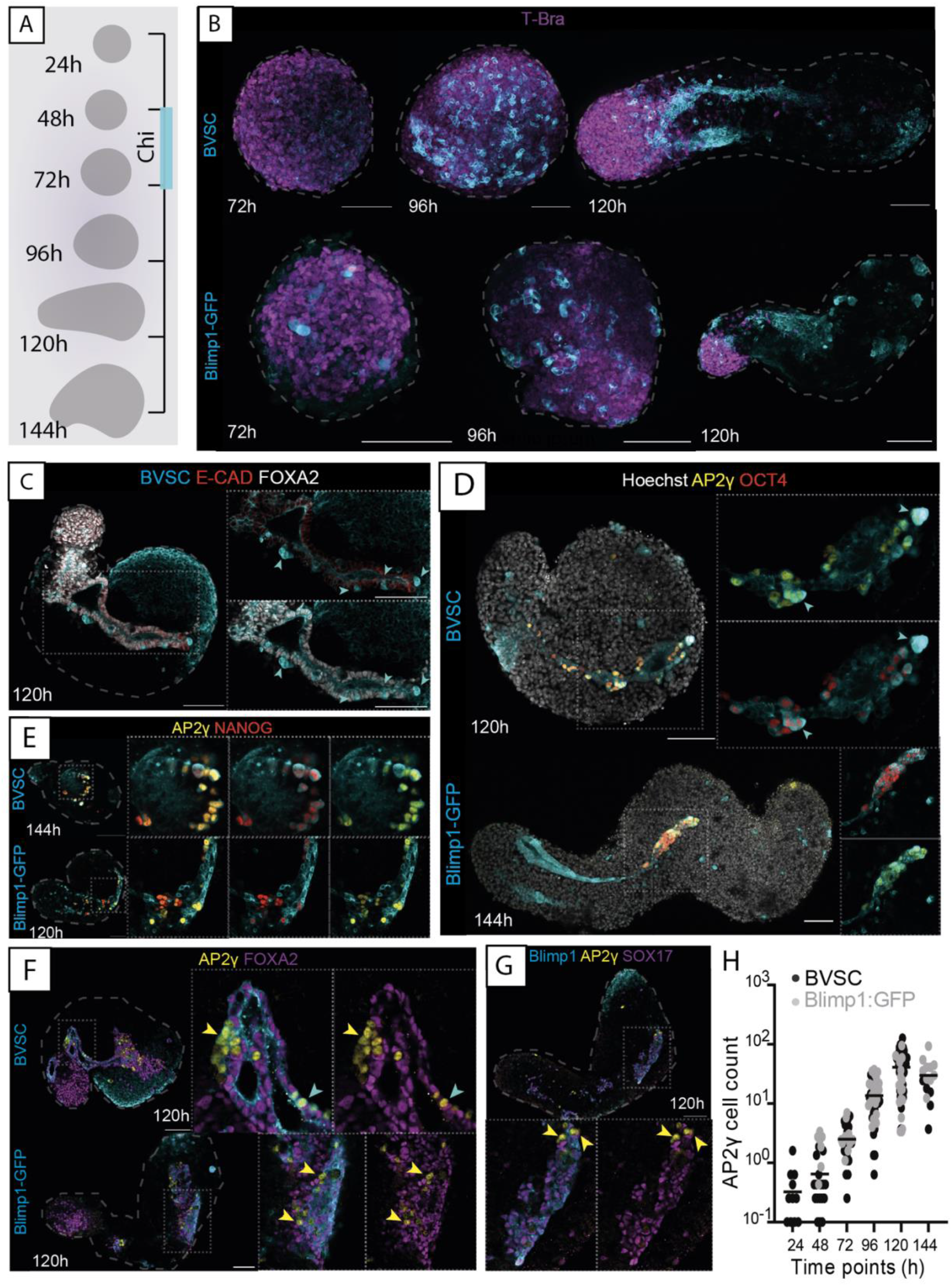
Characterising gastruloid-derived PGCLCs. **(A)** Schematic of gastruloid protocol and morphological changes from 24 to 144h. **(B)** Maximum projection of gastruloids from BVSC and Blimp1-GFP reporter lines. In BVSC gastruloids, Blimp1:mVenus is membrane-targeted while Stella:eCFP is found throughout the cell. **(C)** Z section images of Blimp1-mVenus+ endodermal tracts. **(D)** Expression of AP2γ and OCT4 in gastruloids. **(E)** Expression of AP2γ and OCT4 in gastruloids. **(F-G)** AP2γ-expressing cells do not co-express FOXA2 **(F)** or SOX17 **(G**). **(H)** Cell counts of AP2γ-expressing cells from both Blimp1-GFP and BVSC gastruloids. Black bars represent the mean value at each time point. Cyan arrowheads, *Stella*+ cells; Yellow arrowheads, AP2γ+ cells; Insets, higher magnification images; Dashed line, morphological gastruloid outline from Hoechst staining; Dotted line, magnification region. Scale bars, 100 µm.

We therefore examined the dynamic expression of the PGC-associated gene reporters in these gastruloids. *Blimp1* is expressed in the endoderm of the mouse embryo^5^, and the coalescence of endodermal domains into a tube structure in mouse gastruloids has been previously described^32,42,35^. In our gastruloids, *Blimp1* expression was observed initially in a salt-and-pepper manner across spherical gastruloids at 72h, which then tended to coalesce into domains or clusters of expression in ovoid-shaped gastruloids at 96h (Figure 1B). As mouse gastruloids underwent elongation, the domain of *Blimp1* expression became even more spatially defined, and routinely formed contiguous tracts of *Blimp1* expressing cells running along the anteroposterior axis at 120h (apparent in 78.3% BVSC (n = 60) and 60.8% Blimp1-GFP (n = 74) gastruloids; Figure 1B). These *Blimp1+* tracts also expressed FOXA2, SOX17, E-Cadherin (CDH1) and EpCAM (Supplementary Figure 1B-F), suggesting a definitive endoderm identity. They were internally located and typically formed closed, tube-like structures (Supplementary Figure 1G).

Although the majority of *Blimp1* expressing cells in gastruloids therefore likely represent a definitive endodermal population, we observed several *Blimp1* and *Stella* co-expressing cells that were interspersed within or adjacent to the endoderm tubes in BVSC gastruloids (Figure 1C, Supplementary Figure 1F-G). We reasoned that these were likely to be PGCLCs. Indeed, the PGC marker, AP2γ, was found to be co-expressed with the pluripotency factor OCT4 (also known as POU5F1) and NANOG in a high proportion of these cells (Figure 1D-E, Supplementary Figure 1H) and they did not express endodermal markers, FOXA2 or SOX17 (Figure 1F-G). While *Stella* expression was consistently observed in mouse gastruloids, not all *Stella+* cells were positive for both OCT4 and AP2γ, and often co-expressed only one of these markers (Supplementary Figure 1H-J) suggesting that there might be heterogeneity of *Stella*-expressing cells in Gld-PGCLCs, perhaps related to the temporal range of states observed. Therefore, we also utilised Platelet and Endothelial Cell Adhesion Molecule 1 (PECAM1) expression, which is known to be expressed in PGCs in the mouse embryo (as well as pluripotent and endothelial cells)^43-46^. In 120h gastruloids, PECAM1 was co-expressed in the vast majority of AP2γ (96.65%), OCT4 (95.83%) and *Stella* positive cells (98.3%) in BVSC gastruloids, and when we examined AP2γ expression alongside PECAM1, we observed double positive cells as early as 24h, that were then co-expressed with *Stella* from 72h (Supplementary Figure 2A-C), suggesting that PECAM1 marked the broadest population of Gld-PGCLCs across the time-course. We therefore decided to use both AP2γ and PECAM1, in combination with the endogenous reporters of BVSC or Blimp1-GFP cell lines, as general markers of Gld-PGCLCs.

Gastruloids displayed a consistent and progressive increase in the number of Gld-PGCLCs through the gastruloid timeline from 24h to 120h (Figure 1H). This began as an average 2.42 cells per gastruloid (+/- 2.15 s.d.; 8.3% of gastruloids had no AP2γ expressing cells), which increased to 4.07 +/- 4.16 s.d. at 48h (20% of gastruloids without AP2γ+ cells) and continued to increase to reach a mean average of 90.72 cells per gastruloid by 120h (+/- 49.11 s.d.; 0% of gastruloids had no AP2γ expression, n=57). At 144h the average number of Gld-PGCLCs slightly decreased (71.93 Gld-PGCLCs +/- 37.91 sd; Figure 1H) which mirrored a general decrease in average size of 144h gastruloids (Supplementary Table 1). Likewise, by flow cytometric analysis, a population that was double positive for *Stella-e*CFP and PECAM1 was observed to increase in frequency during BVSC gastruloid development (Supplementary Figure 2C). These estimates of absolute Gld-PGCLC cell numbers are roughly consistent with the equivalent *in vivo* PGC numbers, with approximately ∼100 PGCs found in the E8.5 mouse embryo^47,48^ which represents an equivalent stage to 120h gastruloids^32,33^, and an average doubling time approximating 16.12 hours (Figure 1H), matching the 16 hours estimated for mouse PGCs in the embryo^49^.

#### Dynamic localisation of Gld-PGCLCs

We were particularly interested to note the spatial localisation of the Gld-PGCLCs relative to the endodermal tract, given the role of PGC migration along the endoderm *in vivo*^50,51^. We noted that at 120h, the Gld-PGCLC were often interspersed throughout the endodermal tract along the anteroposterior axis, but by 144h the majority were localised within small clusters of cells at the anterior edge of gastruloids (Figure 2A-B). These each contained an average of 9 cells expressing two or more PGC-associated proteins, and each gastruloid had on average 3.3 clusters (n = 7) (Figure 2C; Supplementary Table 2), similar to PGCs colonising mouse gonads at E10.5^52^. Since Gld-PGCLCs seemed to shift towards the anterior end of the gastruloids relative to the length of the gastruloid (average location at 75.9% +/- 9.95 sd of the gastruloid length starting from the posterior at 144h, n = 15; Figure 2D) we reasoned that they might be moving relative to the axis of maximal elongation of the gastruloid from posterior to anterior (Figure 1E).

**Figure 2:**
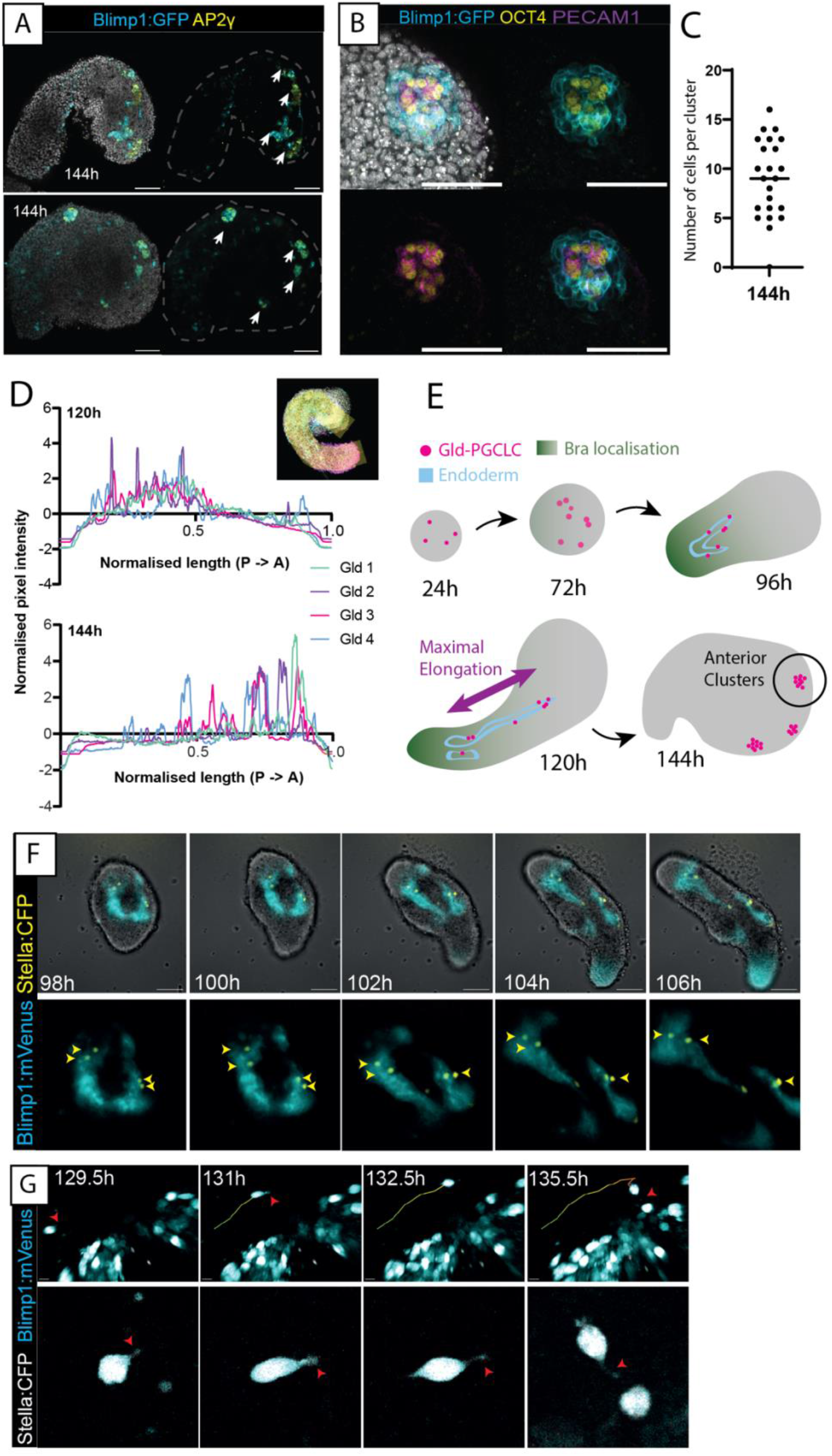
Anterior localisation and movement of Gld-PGCLCs. **(A)** Anterior-localised clusters of AP2γ+ cells at 144h. White arrows, location of discrete clusters. Scale bars, 100 µm. **(B)** High magnification Z slice of an OCT4+ and PECAM1+ cluster at 144h. Scale bars, 100 µm. **(C)** Quantification of the number of cells in each cluster at 144h, as determined by co-expression of at least two of *Blimp1*, AP2γ, PECAM1 or DAZL. Samples from n = 7 gastruloids. Black bar indicates the median average. **(D)** Anteroposterior localisation of AP2γ+ cells along the gastruloid length (see Methods). Gld, Individual gastruloid replicates. Inset, representation of a 120 width line spanning anteroposterior axis of Z max projection gastruloid. **(E)** Schematic representation of Gld-PGCLC localisation within gastruloids across their time-course. **(F)** Widefield time-lapse imaging of a BVSC gastruloid from 98-106h. Top, whole gastruloid image; Bottom, zoom-in of fluorescent reporter domain. Yellow arrowheads, *Stella*+ cells. Scale bars, 100 µm. **(G)** Multiphoton time-lapse images of a BVSC gastruloid from 129.5-135.5h with cell tracking (plotted line). Red arrowheads, cell morphological features associated with active migration. Scale bars, 10 µm.

Indeed, we observed evidence of Gld-PGCLCs cell movement throughout their development in gastruloids. Some of this appears to be due to overall morphological changes associated with gastruloid elongation and might therefore represent a passive relative movement of the Gld-PGCLCs. For instance, Gld-PGCLCs (*Stella*-eCFP expressing) were often already intermingled with endodermal cells (Blimp1-Venus expressing) prior to elongation at 96h (n=29/39 gastruloids) and later became distributed throughout the endodermal tracts concurrent with gastruloid elongation (Figure 2F; Supplementary Movie 1). Since E-cadherin and EpCAM were expressed in both Gld-PGCLCs and the endodermal cells at 96h (Supplementary Figure 3A-B) it is possible that this could potentially mediate the observed close association between the tissues, as has been suggested in the mouse embryo^53,54^, although further experiments would be required to test this hypothesis.

In addition, it is likely that Gld-PGCLCs are also capable of active movement as well as passive relative movement. Using multiphoton microscopy, we observed several instances of *Stella*-positive cells displaying seemingly motile behaviour relative to the gastruloid structure, morphological changes associated with migration including cellular protrusions that appear filopodia-like, and interactions with other *Stella*+ cells (Figure 2G, Supplementary Movie 2). However, such movement was not always strictly posterior-to-anterior, and so the observed shift in relative location of Gld-PGCLCs is likely to be due to both active and passive movement of cells.

Given the apparent role of the endodermal epithelium to coordinate the relative localisation of Gld-PGCLCs to the anterior end of gastruloids, we wanted to investigate the necessity of this endodermal population for Gld-PGCLC localisation. In mouse, *Sox17*-null embryos specify PGCs, but they cannot enter the gut endoderm and are stalled at the hindgut entrance^55^. We therefore generated mouse gastruloids from mESCs that were Sox17^-/-^ (see Methods) or FoxA2^-/-56^ (Figure 3A-L). In both cases, the mutant gastruloids still contained mesoderm and ectoderm, and underwent axial elongation, but the endodermal population was absent, and no epithelial tract was observed. The Gld-PGCLC population was observed at absolute cell numbers equivalent to wildtype gastruloids (Figure 3A), but importantly, they were localised in large clusters at 120h rather than dispersed throughout the length of the gastruloid (Figure 3B, H). This observation strongly supports the notion that the presence of the endodermal tract in gastruloids facilitates the spatially organised movement of Gld-PGCLCs, closely resembling observations in the mouse embryo^55^.

**Figure 3:**
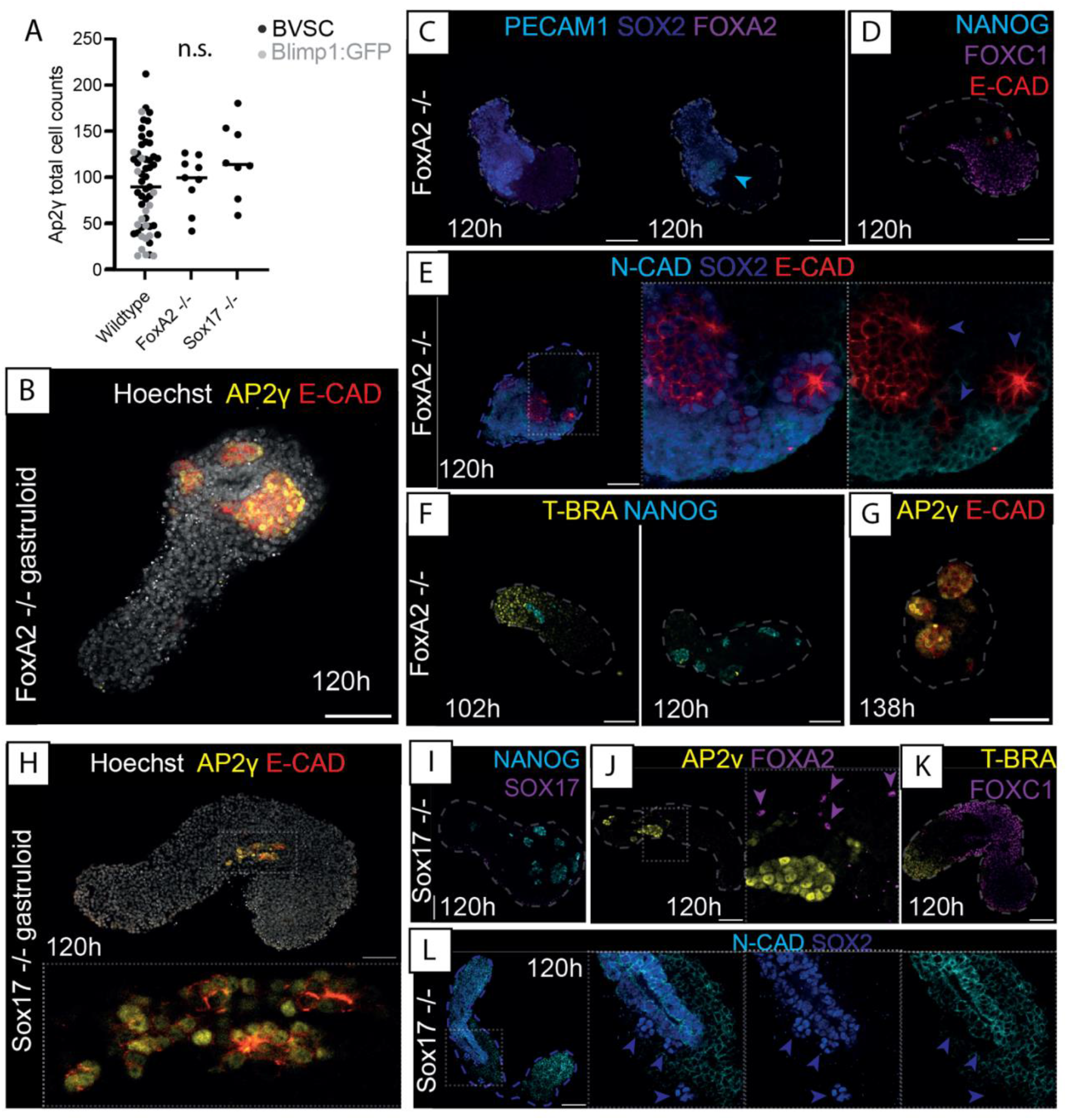
Knockout of Gastruloid Endodermal tissue leads to aberrant Gld-PGCLC localisation but maintains mesoderm and ectodermal populations. **(A)** Quantification of AP2γ+ cell counts in Blimp1-GFP and BVSC gastruloids (Wildtype), alongside FoxA2-/- and Sox17-/- gastruloids. Black bars indicate the median average; n.s., no significant difference. **(B)** AP2γ+ cells localise into large clusters in FoxA2-/- gastruloids and show no E-Cadherin (E-CAD positive) endodermal tracts (AP2γ negative). **(C)** Confirmation of lack of FOXA2 expression detected in FoxA2-/- gastruloids. **(D)** Maintenance of FOXC1 mesoderm in FOXA2-/- gastruloids at 120h. **(E)** Neural ectodermal cell types present in FoxA2-/- gastruloids as evidenced by N-Cadherin (N-CAD) and SOX2 expression. **(F)** Mesodermal T-BRA expression in FoxA2-/- gastruloids at 102h but not 120h. **(G)** Later stage putative Gld-PGCLC in 138h FoxA2-/- gastruloids. **(H)** AP2γ+ cells localise into large clusters in Sox17-/- gastruloids and show no E-Cadherin (E-Cad positive) endodermal tracts (AP2γ negative). **(I)** Confirmation of lack of SOX17 expression detected in SOX17-/- gastruloids. **(J)** Presence of several scattered FOXA2+ cells (purple arrowheads) in SOX17-/- gastruloids. **(K)** Maintenance of FOXC1 mesoderm in SOX17-/- gastruloids at 120h. **(L)** Neural ectodermal cell types present in SOX17-/- gastruloids as evidenced by N-Cadherin (N-Cad) and SOX2 expression. Blue arrowheads, SOX2+, N-Cad- cells likely to be Gld-PGCLCs. **(A-L)** Insets, higher magnification images; Dashed line, morphological gastruloid outline from Hoechst staining; Dotted line, magnification region. Scale bars, 100 µm.

#### Maturation of Gld-PGCLCs

The morphological clustering of Gld-PGCLCs in the anterior of 144h gastruloids was highly reminiscent of gonadal germ cell clusters found in the mouse embryo at E11.5^10,52^. We therefore wondered whether these anterior-localised Gld-PGCLCs were undergoing further maturation, particularly in the form of epigenetic remodelling. Indeed, we observed that the histone modification, H3K27me3, which has been shown to be associated with PGC maturation to a germ cell fate^57,58^, was co-localised with AP2γ at 144h (35% co-expression; Figure 4A-B). Similarly, the DNA modification mark, 5hmC, was also co-localised with Gld-PGCLCs in anterior clusters of cells in 144h gastruloids (45% co-expression; Figure 4C-D), another hallmark of PGC maturation^59,60^. Since DNA demethylation is required to de-repress the promoter of the germ cell gene *Dazl*, which itself is required to facilitate the maturation of germ cells towards sex-specific stages in a process called ‘licensing’^12^, we examined the expression of DAZL in gastruloids. Surprisingly, we observed clear DAZL protein expression in Gld-PGCLCs at 120h (mean = 28 +/- 15.46 s.d. cells per gastruloid, n = 8) which stayed consistent in 144h gastruloids (mean of 46.6 +/- 47.93 s.d. cells per gastruloid, n = 15; Figure 4E) and were localised particularly in anterior clusters (Figure 4F). Furthermore, the DAZL was co-expressed with AP2γ (21% co-expression; Figure 4G) and we generally found DAZL expression in cells that had lower levels of NANOG expression (Figure 4H), potentially relating to its role in downregulating pluripotency factors during germ cell maturation^61^. As such, it seems that the Gld-PGCLCs begin to undergo a maturation process that, to some extent, mirrors the post-migratory/gonadal stage development of PGCs *in vivo,* and which might be directly mediated by their local environment.

**Figure 4:**
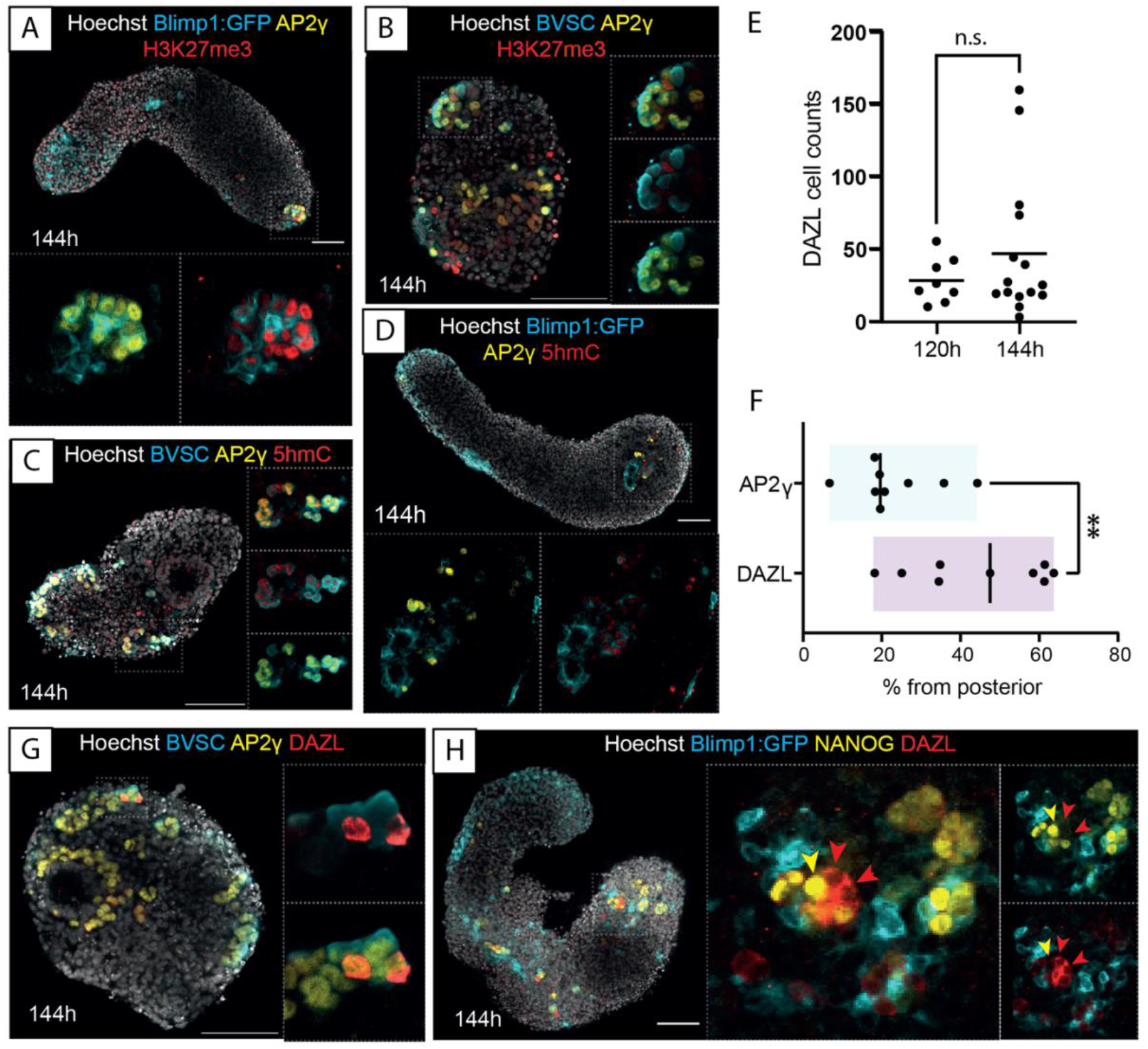
Maturation of Gld-PGCLCs in epigenetic and protein expression changes associated with germ cell determination. **(A-B)** Histone H4 trimethylation of K27 (H3K27me3) in Gld-PGCLCs in Blimp1-GFP **(A)** and BVSC **(B)** gastruloids at 144h. In BVSC gastruloids, Blimp1:mVenus is membrane-targeted while Stella:eCFP is found throughout the cell. **(C-D)** 5-Hydroxymethylcytosine (5hmC) in Gld-PGCLCs in Blimp1-GFP **(C)** and BVSC **(D)** gastruloids at 144h. **(E)** Quantification of DAZL-expressing cells in BVSC and Blimp1-GFP gastruloids. Black line represents the mean cell count. n.s., no significant differences. **(F)** Quantification of Gld-PGCLC localisation along the anteroposterior axis, using the posterior-most detected expression from each gastruloid as a percentage of total length (see Methods for details). Black line represents the median value. **(G-H)** DAZL expression in 144h BVSC **(G)** and Blimp1-GFP **(H)** gastruloids. Yellow arrowhead, NANOG+, DAZL-cell; Red arrowhead, NANOG-, DAZL+ cells. Insets, higher magnification images; Dashed line, morphological gastruloid outline from Hoechst staining; Dotted line, magnification region. Scale bars, 100 µm.

We hypothesised that local signalling or niche properties of surrounding cells in the anterior region of the gastruloid could be supporting these cell clusters. Indeed, we frequently observed high-level expression of GATA4 in several cells near the Gld-PGCLC clusters (Supplementary Figure 4A-C). In support of this observation, closer examination of extant spatial transcriptomics datasets from 120h mouse gastruloids^33^ showed an anterior localisation of *Gata4*, an early marker of the developing bipotent gonad^62^ and *Cxcl12* (also known as *Sdf1*), a chemokine thought to be responsible for directional migration in the mouse embryo^63,64^ (Supplementary Figure 4D). It is possible that these spatially-localised supporting cells enable the maturation of Gld-PGCLCs to post-migratory stages of development, as they begin to express not only DAZL but also GCNA1, a marker of post-migratory PGCs *in vivo^65^* (Supplementary Figure 4E).

### Transcriptomic Gld-PGCLC characterisation

Given the general signature of PGC-identity observed in Gld-PGCLCs, including the surprisingly mature status of DAZL- and GCNA1-expression, we wanted to compare our Gld-PGCLCs to known populations of PGCs, both *in vivo* and *in vitro,* at the transcriptomic level. To do this, we sorted Blimp1:mVenus+, SSEA1+ cells, PECAM1+ cells and Stella:eCFP+ cells from 120h gastruloids and performed 10x single-cell RNA-sequencing (Methods; Supplementary Figure 5A). Once integrated into a single 120h dataset, we identified 8 distinct clusters of cell identities (clusters 0 to 7), including 5 that we denoted to be putative PGCLCs due to expression of genes including *Dppa3/4/5A, Nanog, Oct4 (Pou5f1), Sox2, Blimp1 (Prdm1)* and *Ap2*γ (*Tfap2c*; Supplementary Figure 5B). In addition, some cells within these clusters also expressed genes including *Dazl*, *Ddx4* and *Tex14* which are known markers of later stage PGCs in the mouse embryo. While each sorted population contributed to these PGC-like clusters, we also noted additional populations including a putative endoderm-like population (Cluster 6), endothelium (Cluster 7) and mesoderm, including somitic cell types (Cluster 5) that were apparent in our data (Supplementary Figure 5B, C). To further confirm that our sorting strategies were indeed capturing the population of Gld-PGCLCs we compared our data to extant mouse gastruloid scRNA-seq data^33^, and confirmed a high degree of concordance between both PGCLC populations (Supplementary Figure 5D-E). We therefore filtered our cells using the previously defined PGCLC population from mouse gastruloid scRNA-sequencing data^33^ for all downstream analysis.

One of our major questions was whether these cells were equivalent to *in vivo* PGC cell types, and if so, which developmental timepoint was best matched by the *in vitro* Gld-PGCLCs. To assess this, we projected our Gld-PGCLCs onto a well-characterised and extensive map of *in vivo* germ cell development between E6.5 and adulthood (8-10 weeks) at 28 sampled timepoints from Zhao and colleagues^66^. Surprisingly, we found a very close match between our Gld-PGCLC cells and *in vivo* PGCs at the mitotic and mitotic arrest PGC stage of development which were found in E13.5-15.5 stage embryos (Figure 5A-D). This is particularly remarkable given that traditional embryoid body-derived PGCLCs are thought to stall at E9.5-E10.5 stages^15^. We therefore directly compared our Gld-PGCLC dataset to a published single cell dataset from EB-derived PGCLCs at day 6^29^ with the *in vivo* PGC dataset. We found that the EB-PGCLCs were relatively heterogenous, and their projection spanned cell types from specification PGCs, to migrating PGCs and as late as mitotic PGCs (E8.5 to E13.5) while our Gld-PGCLCs were more homogeneous and clearly more advanced on the projection, and approximated particularly mitotic arrest PGCs (E13.5 to E15.5; Figure 5E-G).

**Figure 5:**
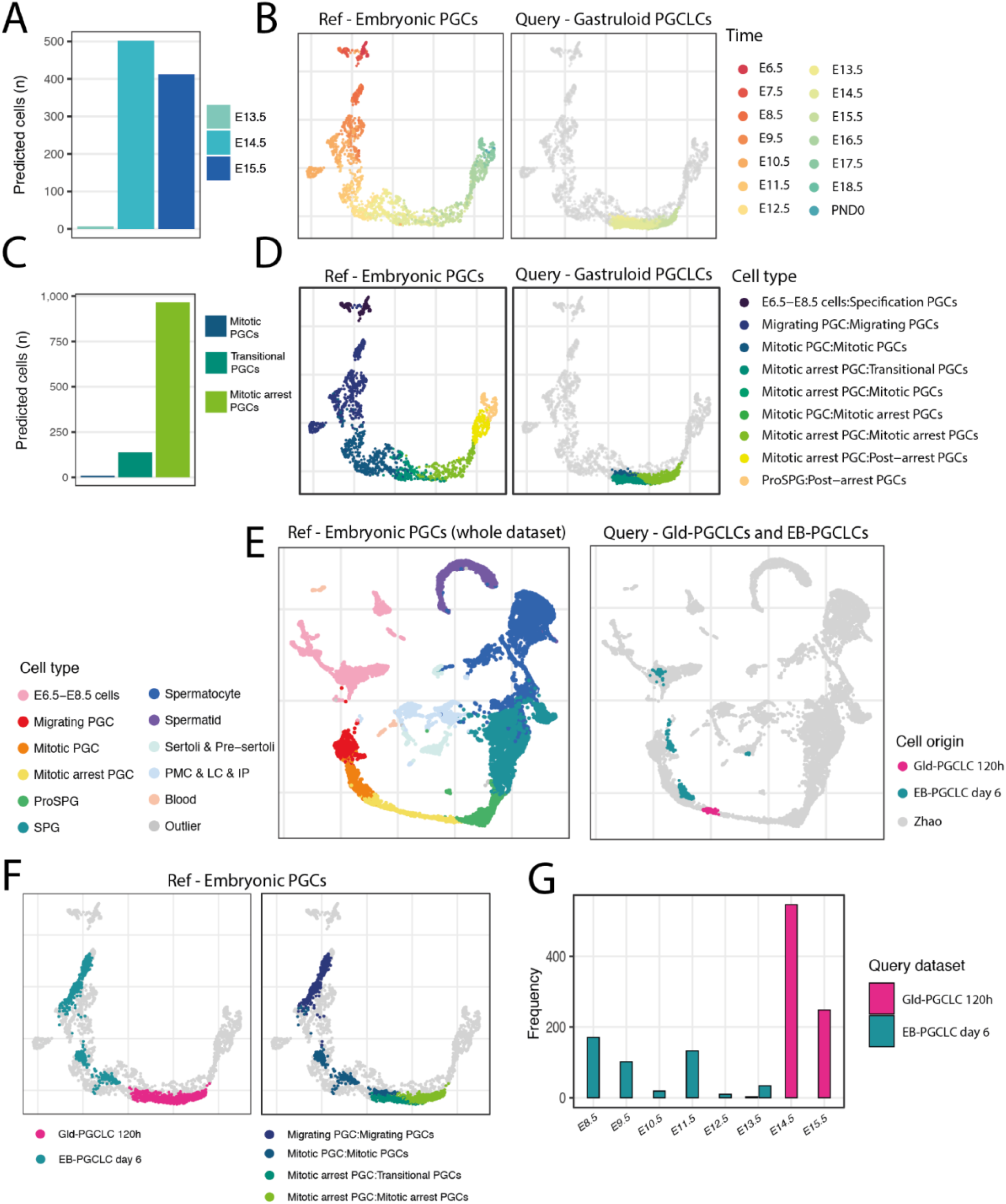
Single-cell transcriptomic comparison between Gld-PGCLCs and an *in vivo* PGC dataset^66^. **(A, C)** Quantification of label transfer prediction from Gld-PGCLCs (0.6+ max prediction score) in terms of embryonic time **(A)** and cell stage **(C). (B, D)** UMAP of PGC-only cell types from Zhao and colleagues, in terms of time **(B)** and stage **(D) with Gld-PGCLC embedded. (E)** UMAP projection of Gld-PGCLC (0.9+ max prediction score) and published EB-PGCLCs^29^ into the *in vivo* UMAP of the full dataset. **(F)** Comparison of UMAP projection of Gld-PGCLCs and published EB-PGCLCs onto in vivo PGC dataset, by origin (left) and cell state (right). **(G)** Frequency of cell transfer labels from EB-PGCLCs or Gld-PGCLCs (0.6+ max prediction score) onto the *in vivo* PGC dataset, by embryonic time point.

Together, this transcriptomic analysis of Gld-PGCLCs alongside the observation of protein level DAZL expression, epigenetic remodelling and cell morphological behaviours suggests that gastruloids might enable the development of more mature PGC-like states *in vitro*, without the need for additional gonadal co-culture.

### Endogenous signalling control of Gld-PGCLC specification

Since no exogeneous manipulation of the gastruloids was performed that might particularly bias towards a germ cell-like fate, we hypothesised that Gld-PGCLC specification and maturation must be coordinated by local, self-organised signalling feedback mechanisms between populations of cells present in the gastruloid. We therefore sought to manipulate the endogenous signalling environment of gastruloids and examine the resultant effect on the Gld-PGCLC population to better understand how these endogenous signals were acting.

We initially focussed on BMP signalling pathway, since it has been reported to be required for PGC specification *in vivo*^8,67-70^ and *in vitro*^71,7^, although this has been brought into question by recent reports^72,73^. Surprisingly, we found that addition of BMP4 ligand did not lead to any significant increase in Gld-PGCLC numbers compared to control gastruloids (Figure 6A-B, Supplementary Figure 6A-B). Likewise, no co-localisation of phosphorylated SMAD1/5/8 (pSMAD1/5/8) was found in Ap2γ cells at 24h in BVSC or 48h in Blimp1-GFP gastruloids (Supplementary Figure 6C-D) and in general, very little pSMAD1/5/8 was detected in the gastruloids until 96h, where the distribution was polarised towards the anterior pole but was never observed to co-localise with AP2γ (Supplementary Figure 6C-D). This is consistent with spatial transcriptomics data that reported an anterior bias of BMP signalling in gastruloids from late stages^39,74^ but implies that downstream BMP signalling might not be active in the Gld-PGCLCs themselves. Indeed, addition of the BMP inhibitor, Dorsomorphin homolog 1 (herein, DMH1; a selective inhibitor of ALK2), to gastruloids from 24 to 48h did not produce discernible morphological differences in axial elongation when compared to the DMSO control, and both contained AP2γ/*Stella* positive cells (Figure 6C). However, a significant increase was found in absolute AP2γ cell count (p=0.0003; 43 +/- 22.52 sd mean cells per gastruloid) and proportion relative to gastruloid volume (p=0.0191, 5.78 +/- 4.2 sd mean cells per gastruloid) in gastruloids exposed to DMH1 (Figure 6C-D). Consistent with this, higher concentrations of DMH1 resulted in further significant increases in AP2γ+ cell count (Figure 6C, Supplementary Figure 7A-C), and LDN 193189 (herein, LDN; an ALK3 inhibitor) treatment likewise did not inhibit Gld-PGCLC formation (Supplementary Figure 7A).

**Figure 6:**
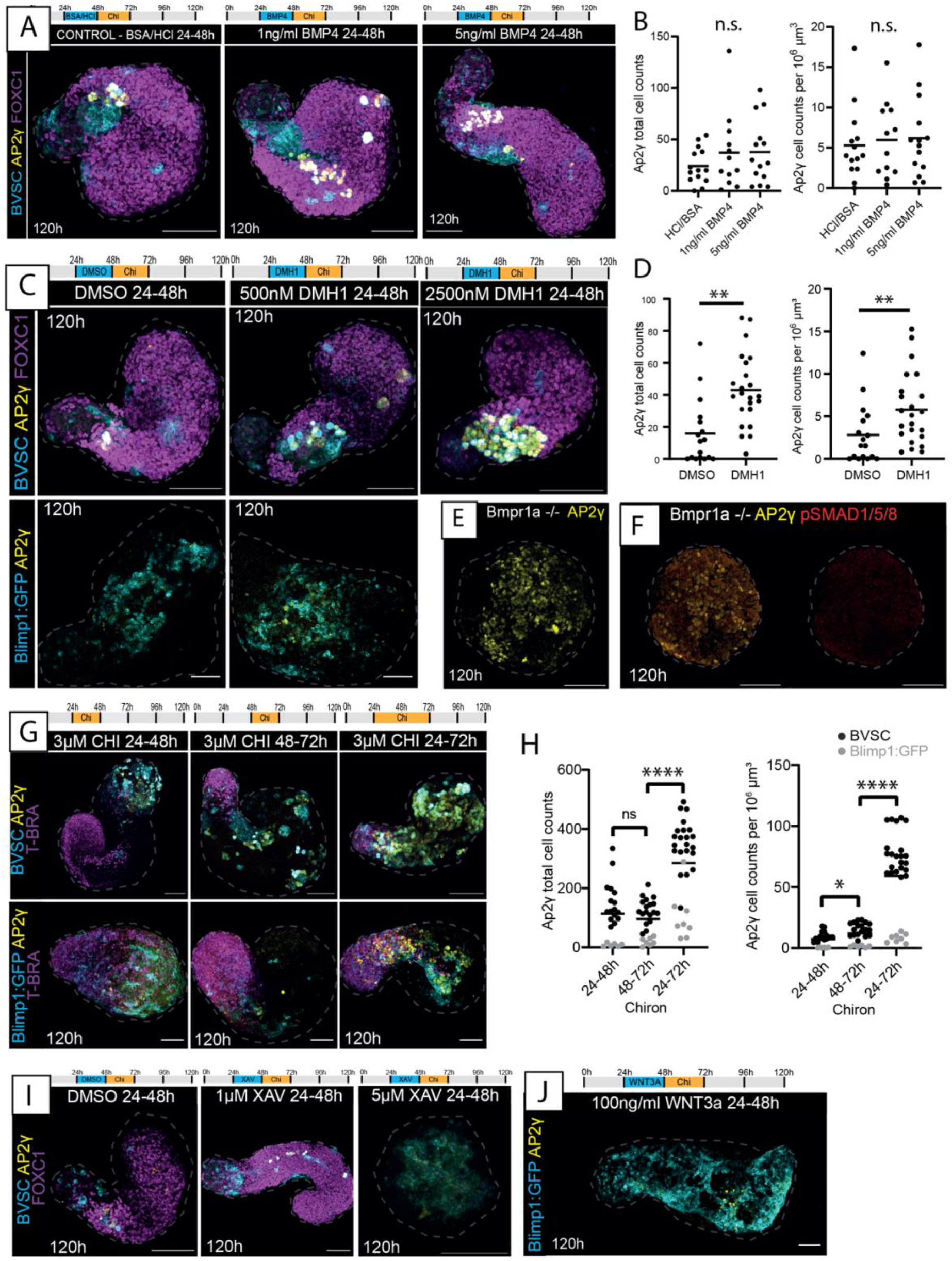
BMP and Wnt signalling modulation in Gld-PGCLCs. **(A)** Maximum projection images of BVSC gastruloids following BMP application at timepoint and concentrations indicated. In BVSC gastruloids, Blimp1:mVenus is membrane-targeted while Stella:eCFP is found throughout the cell. **(B)** Quantification of AP2γ+ cells in conditions indicated from BVSC and Blimp1-GFP gastruloids at 120h. n.s., no significant differences. **(C)** Maximum projection images of gastruloids following BMP inhibition by DMH1 application at timepoint and concentrations indicated. **(D)** Quantification of AP2γ+ cells in DMSO or 500nM DMH1, from BVSC and Blimp1-GFP gastruloids at 120h. **(E)** Gastruloids made from BMPR1A -/- cell line, showing aberrant gastruloid morphology with lack of elongation, and significant numbers of AP2γ+ cells. **(F)** Absence of pSMAD1/5/8 in BMPR1A-/- gastruloids at 120h. **(G)** Maximum projection of gastruloids exposed to different timing of Chi application, as indicated. **(H)** Quantification of AP2γ+ cells in conditions indicated, from BVSC and Blimp1-GFP gastruloids at 120h. **(I)** Maximum projection of BVSC gastruloids exposed to Wnt signalling inhibition by application of XAV. **(J)** Maximum projection of Blimp1-GFP gastruloid exposed to WNT3a at timepoint shown. **(B, D, H)** Black bars represent the mean average. **(A-J)** CHI, CHIR99021; XAV, XAV939. Dashed line, morphological gastruloid outline from Hoechst staining; Scale bars, 100 µm.

To further explore this surprising relationship between BMP signalling and Gld-PGCLC specification, we generated gastruloids from BMPR1a null mESCs^75^. These gastruloids did not elongate (Figure 6E-F), perhaps consistent with the reported reduced Nodal/Activin signalling found in *Bmpr1a* null embryos^75^ and the requirement for Nodal signalling in symmetry breaking and elongation in gastruloids^39^. However, they did show evidence of differentiation towards endoderm, mesoderm, and ectodermal populations (Supplementary Figure 7D-I). Surprisingly, they also contained AP2γ expressing cells (Figure 6E, Supplementary Figure 7D) in significantly higher proportions than observed in non-mutant Blimp1-GFP/BVSC gastruloids (Supplementary Figure 7D). Together, these results suggest that BMP signalling is not strictly required for PGCLC specification in the gastruloid model, and indeed may even have a repressive effect on the Gld-PGCLC fate, at least at the timepoints assessed here.

We then turned our attention to the Wnt signalling pathway, which has also been proposed to support PGC specification *in vivo^7,17^*. Since the standard gastruloid protocol includes a 24h pulse of CHIR-99021 (herein, Chi; an inhibitor of GSK3β), between 48-72h post aggregation, we decided to modulate the time interval of addition of Chi to examine the effect on Gld-PGCLCs. Moving the Chi addition 24h earlier altered gastruloid morphology but did not inhibit the presence of AP2γ cells (Figure 6G). However, extending the Chi exposure to between 24-72h post-aggregation resulted in significant (p < 0.0001) increase in AP2γ positive cells in both absolute (mean 285.4 +/- 138.1 sd cells) and relative (mean 59.13 +/- 34.39 sd) cell numbers, although we noted a line specific difference between the Blimp1-GFP and BVSC lines (Figure 6G-H). The increase in AP2γ cells was specific to this time window, as Chi treatment for an equivalently prolonged period of 48h between 0-48h post-aggregation in the BVSC gastruloids did not significantly alter the AP2γ cell number (p = 0.43) even though it did result in clear morphological changes (Supplementary Figure 8A) and later addition of Chi (72-96h) led to a significant decrease in Gld-PGCLCs (Supplementary Figure 8B-C). However, although changing the timing of Chi exposure had an obvious effect on Gld-PGCLC numbers, altering the concentration of Chi between 48-72h did not change the number of AP2γ+ cells relative to the total gastruloid (Supplementary Figure 8D-G). Together, these results suggest that gastruloid PGCLCs are sensitive to Wnt signalling modulation, but that this occurs within a specific temporal window, in a time-dependent but not concentration-dependent manner.

In addition, it is likely that endogenous as well as exogeneous Wnt signalling may be driving PGCLC formation in gastruloids. Gastruloids without a Chi pulse still contained AP2γ expressing cells (Supplementary Figure 8H-I) but BVSC gastruloids exposed to Wnt inhibition by addition of XAV393 (XAV) resulted in loss of Gld-PGCLCs (p = 0.033, Figure 6I, Supplementary Figure 8I). Additionally, the supplementation of 500ng/ml WNT3A led to a significant (p = 0.0023) increase in Gld-PGCLCs in the BVSC gastruloids (Supplementary Figure 8H,I) although addition of 100ng/ml WNT3a on Blimp1-GFP gastruloids did not lead to significant changes (Figure 6J). Taken together, these observations suggest that Wnt signalling is indeed necessary for the specification of Gld-PGCLCs, and that gastruloids are particularly sensitive to the effect of this pathway between 24-72h post-aggregation.

Finally, we turned our attention to the FGF signalling pathway, as an *in vitro* study found that FGF inhibition during mesodermal induction resulted in the formation of mouse PGCLCs^76^. Phosphorylated ERK (pERK) was observed sporadically with no discernible spatial polarisation in 24/48h gastruloids, and neither was it specifically associated with AP2γ positive cells (Supplementary Figure 9A). Perturbation of the FGF pathway by addition of the FGF signalling inhibitor, PD0325901 (herein, PD03), between 24-48h resulted in a marked increase in AP2γ expressing cells accompanied by loss of gastruloid elongation and disruption of FOXC1, a marker of anterior mesoderm (Figure 7A). The AP2γ-expressing total cell count was significantly (p < 0.0001) higher than the DMSO control (mean average of 290.3 cells +/- 95.45 sd) and increased in a concentration-dependent manner when adjusted for gastruloid volume (Figure 7A-B, Supplementary Figure 9B-C). This observation was independent of Chi, as an increase in AP2γ expression was also observed when PD03 was added to gastruloids in the absence of the Chi pulse (Supplementary Figure 9B-C). The FGF inhibition-induced increase in AP2γ was also timeframe specific, with the largest change in AP2γ number following PD03 addition between 24-48h (Supplementary Figure 9D).

**Figure 7:**
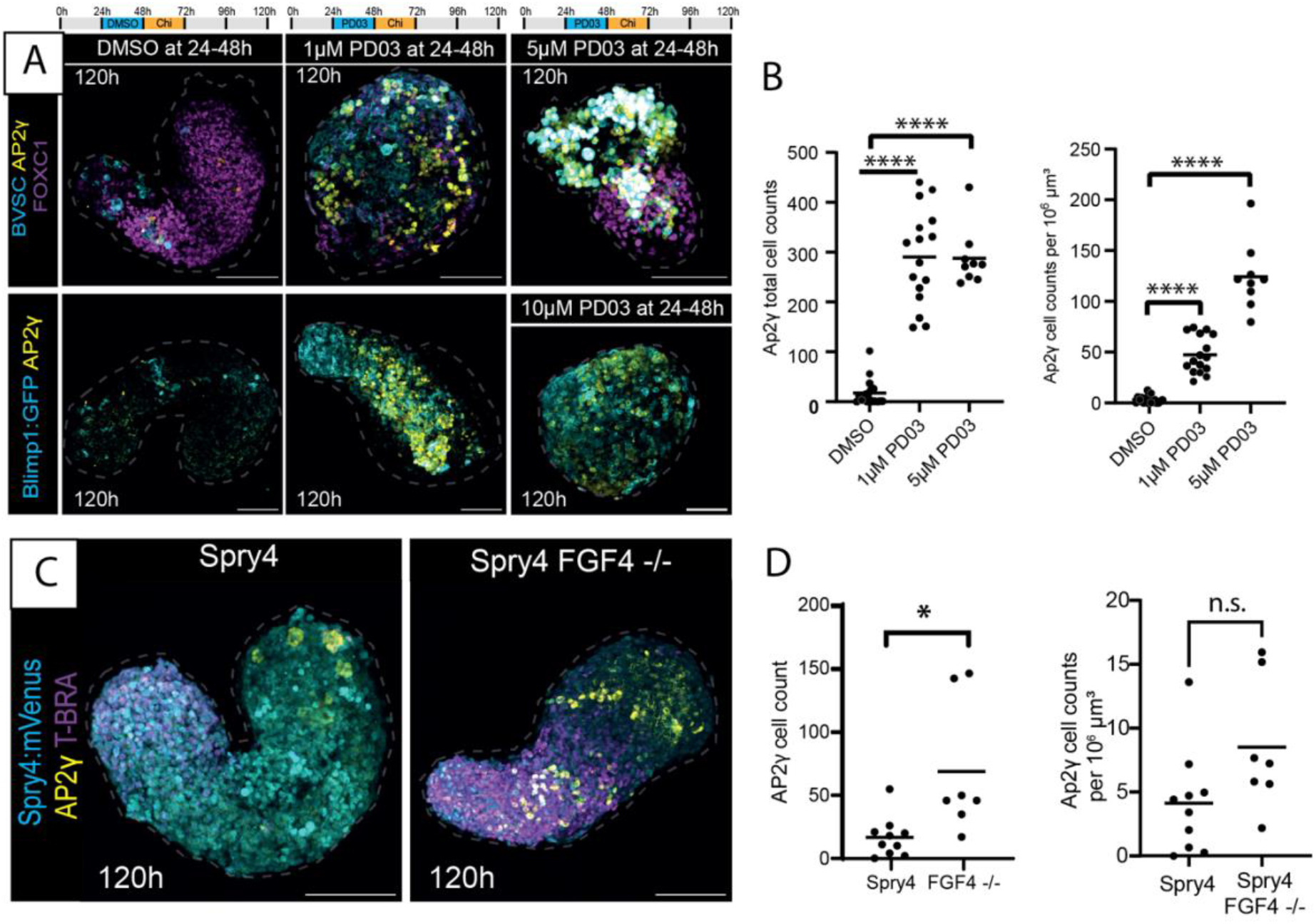
FGF signalling modulation in Gld-PGCLCs. **(A)** Maximum projection of gastruloids exposed to FGF signalling inhibition through PD0325901 (PD03). In BVSC gastruloids, Blimp1:mVenus is membrane-targeted while Stella:eCFP is found throughout the cell. **(B)** Quantification of AP2γ+ cells in conditions indicated, from BVSC and Blimp1-GFP gastruloids at 120h. **(C)** Maximum projection of Spry4:mVenus FGF4 -/- gastruloids at 120h. **(D)** Quantification of AP2γ+ cells in non-mutant Spry4:Venus gastruloids and in Spry4:mVenus FGF4 -/- gastruloids at 120h. n.s., no significant differences. Dashed line, morphological gastruloid outline from Hoechst staining; Scale bars, 100 µm.

To further explore the role of FGF signalling on Gld-PGCLC specification, we made gastruloids from cells containing a fluorescent reporter of the downstream target of the FGF pathway, *Spry4* (Spry4-Venus), as well as this same reporter line with FGF4 knock-out (*Spry4*-Venus; FGF4^-/-^)^77^. *Spry4*-Venus expression was found to be biased towards the more posterior end of the gastruloids, consistent with a posterior FGF signalling gradient in gastruloids^39^ and in the gastrulation-stage embryo^78,79^ and, like our pERK stainings, *Spry4* reporter expression did not overlap specifically with the AP2γ population (Figure 7C). However, gastruloids generated from the FGF4 mutant cells had a significant increase in AP2γ positive cells (p = 0.0398; Figure 7D-E) akin to our results with small molecule inhibition of this pathway. While we cannot rule out that these FGF-modulated AP2γ positive cells show differences to Gld-PGCLCs in the absence of exogenous FGF signalling, these results are suggestive of potential Gld-PGCLC sensitivity to FGF signalling levels that should be investigated in future studies. Together, these signalling modulation experiments suggest that there are specific time-windows that are sensitive to signalling pathway perturbation in mouse gastruloids, that might correspond to times at which cellular populations undergo cell fate decisions or emerge as new cell types, and particularly implicate the Wnt and FGF pathways as key modulators of Gld-PGCLCs in gastruloids.

## Discussion

We have shown that gastruloids generated from established PGC reporter lines contain a population of cells that display key features of PGCs, including co-expression of pluripotency and PGC-associated markers, that we call Gld-PGCLCs. Our findings, combined with those by others^42,33-35,41^, demonstrate that Gld-PGCLCs appear to be general feature present in mouse gastruloids, despite the fact that gastruloids self-organise in the absence of extraembryonic tissues^39^. Our results have shown that gastruloids are able to specify a population of PGC-like cells and support the continued maturation of this population towards late-PGC identities, dynamically recapitulating many aspects of their *in vivo* counterparts in gene/protein expression, epigenetic changes, and cell behaviour.

In addition, the Gld-PGCLCs generated here show advanced maturation equivalent to ∼E14.5 stage *in vivo* PGC development, that far surpasses traditional EB-PGCLC approaches that are believed to stall at approximately E9.5-10.5 stages^15^. One example of this is in the expression of DAZL, a late germ call marker that is required for germ cell determination^12,13^, which Gld-PGCLCs express at 120h but is not typically reached in EB-PGCLCs^15^, except in the presence of additional expansion factors such as forskolin and rolipram^80^. It is likely that the close association of Gld-PGCLCs with neighbouring tissues in gastruloids, including the early primitive streak-like domain, the epithelial endodermal tract and the GATA4+ anterior niche cells, strongly support the notion that mouse gastruloids benefit from organised co-development of Gld-PGCLCs alongside somatic populations. Potentially, this could explain their apparent maturation, as local endogenous signalling alongside dynamic cell movements might be optimising the developmental time-course of these cells towards developmentally-faithful fates^14^. However, it is indeed surprising that Gld-PGCLCs are able to reach states equivalent to embryonic E14.5 PGCs by 144h, given that previous studies have suggested that gastruloids at this experimental timepoint are overall most similar to ∼E9.5 stages^32^. It is possible that this observation therefore reveals potential intrinsic properties of PGC(LC)s that, in the embryo, need to traverse long distances to reach the incipient gonads, but may already be competent to reach mitotic arrest stages given the right environment in a simplified *in vitro* system. However, further studies would be required to test this hypothesis.

Our perturbation experiments likewise challenge the role of different signalling pathways in mouse PGC specification. While BMP signalling has been proposed to principally mediate initial specification of the PGC lineage^8,7^, we find little evidence that BMP is required for Gld-PGCLC specification. These results are directly comparable to those performed by Morgani and colleagues^73^ who similarly showed that PGCLCs can be induced in BMPR1A-/- embryoid bodies, and indeed that the proportion of AP2γ+ PGCLCs increases in this case. Together, such results are challenging the notion that BMP signalling is directly required for PGC(LC) induction. Instead, Wnt and FGF signalling appear to be playing a greater role in determining the germline-to-soma balance of cell type proportions in gastruloids. It is possible that BMP signalling is a required feature of mouse embryonic PGC specification primarily because of its role in the extraembryonic-to-embryonic signalling cascades that are necessary to localise the site of presumptive PGC specification to the Proximal Posterior Epiblast^1,81-83^. In gastruloids lacking extraembryonic tissues, the competence of the cells to form PGCLCs is likely to be global rather than localised, similar to experiments isolating epiblast from visceral endoderm and extraembryonic ectodermal tissues^82,7^. However, unlike those early epiblast isolation assays, in this case the time-window of competence appears to have shifted beyond the BMP-receptive stage to a Wnt-receptive stage, particularly between 24-72h of the gastruloid protocol, consistent with similar timepoints in the mouse embryo, at about E5.75 to E6.75^7,84,17^. After this, FGF may well act to ‘fine-tune’ the number and balance proportions of PGCLCs, as has been shown across early cell fate decisions^85^, and similar to its function in separating PGCs from the soma in the Axolotl^86^. Whether this observation is partly specific to the *in vitro* gastruloid context or reflects a more general feature of mouse PGC specification and regulatory control remains to be seen.

Future research may help to unravel further the signalling mechanisms at play within such systems, including the cross-talk between signalling pathways, and the relationship between tissue types and signalling dependencies, potentially leading to answers to longstanding questions that still exist such as how PGCs form in the PPE along with multiple other cell types exposed to the same signalling environment and what exactly determines the cell proportions^14^. In addition, gastruloids have been more recently been generated from human PSCs^74^, so it would be very interesting to see whether these findings translate into human gastruloids, particularly given the current debate about the epiblast or amniotic origin of PGCs in the human embryo^87^.

Overall, our observations highlight the experimental tractability of *in vitro* embryo-like models to generate rare cell types within a native embryo-like context that opens a new route towards exploring exactly how tissue and cell interactions might mediate cell fate specification in embryogenesis. In addition, the Gld-PGCLCs generated here represent an advanced maturation state that has not previously been achieved *in vitro* without the exogeneous application of PGC-specific maturation factors or gonadal co-culture. Both of these features; their maturity and their inherent co-development; represent a unique advantage of using embryo-like model systems over traditional directed differentiation or disorganised EB systems, since cell types are specified in a manner that harnesses the mechanisms that are used by the embryo itself.

## Material and Methods

### Cell culture and maintenance

The following mESC lines were used: Blimp1-GFP^4^ (kindly provided by A. Surani), Blimp1:mVenus Stella:eCFP (BVSC)^88^ (kindly provided by M. Saitou), Sox17 -/- (as described below), FoxA2 -/-^56^ (kindly provided by H. Likert), BMPR1a -/-^75^ (kindly provided by T. Rodriguez), Spry4:Venus and Spry4:Venus FGF4 -/-^77^ (kindly provided by C Schroeter). All mESC lines were cultured in 2iLif in N2B27 (NDiff227 Takara Bio, Y40002, supplemented with 3µM CHIR99021 (Chi), 1µM PD0325901 (PD03) and 11ng/ml mLIF) on gelatinised (0.1% gelatin) tissue culture flasks or 6-well plates kept in humidified incubators at 37°C, 5% CO_2_. Cells were passaged into new flasks or plates every two days with media exchanged daily.

### Generation of Sox17 -/- cell line

Cells were grown for at least two passages prior to transfection. Cas9/gRNA targeting was used to generate strand breaks alongside homologous recombination with a targeting vector^89^. An eGFP sequence was knocked-in to both alleles of the *Sox17* gene by plasmid transfection. Guide RNAs (gRNAs) were designed to target PAM sequences at the start and end of the protein coding sequence (Table 1). gRNAs were ligated into the PX459-Cas9 plasmid^90^ after cleavage with BbsI. The correct integration of the gRNAs was confirmed after cloning by Sanger sequencing using the hU6-F oligonucleotide (see Table 1). Cells were transfected with three plasmids (*Sox17 GFP*, *PX459-gRNA1*, *PX459-gRNA2*) by incubation with FuGene HD (Promega, E2311) following a previously described protocol^91^. Transfected cells were grown under selection with puromycin (Thermo Fisher, A1113803) and clones were picked for expansion. Genomic DNA was prepared from the primary clones for genotyping by PCR, with the primers as described in Table 1.

**Table 1:**
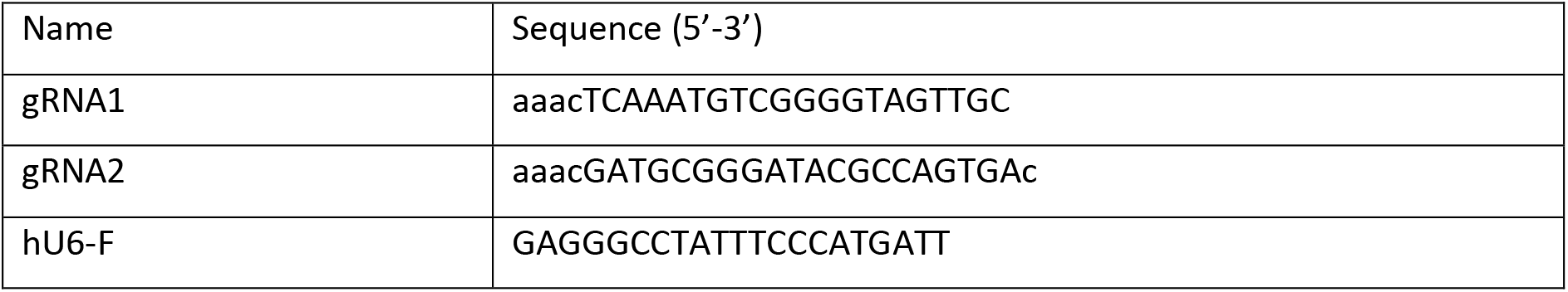
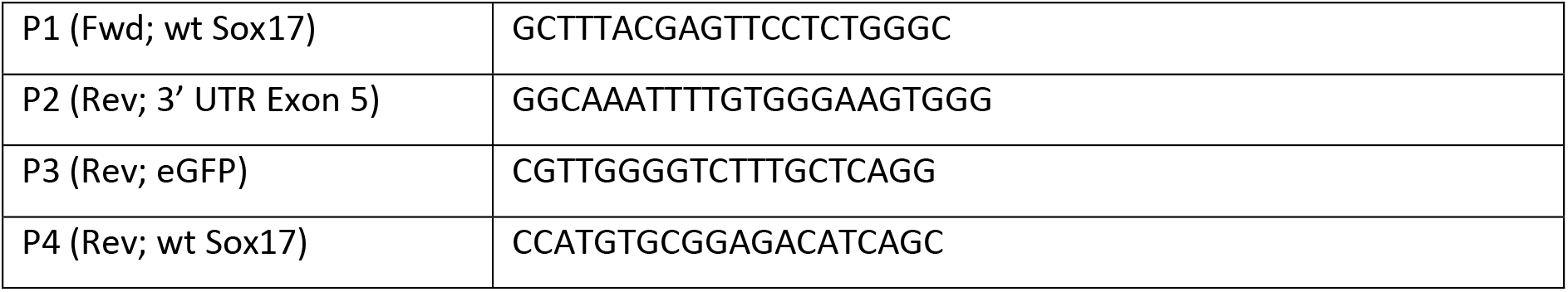
Guide RNA sequences for CRISPR/Cas9 targeting and validation.

### Gastruloid generation

Gastruloids were prepared following the previously reported protocol^92^. Briefly, mESCs were trypsinized and pelleted, with the cell pellet washed in PBS before repeating the process then resuspending in N2B27. The cells were counted and diluted to provide 300 cells per well, before pipetting into U-bottom suspension 96-well plates (Greiner), except in the case of BVSC which were pipetted into cell-repellent, ultra-low attachment 96-well plates (Greiner). Aggregates were incubated at 37°C, 5% CO_2_ in a humidified incubator. After 48 hours, N2B27 supplemented with 3µM Chi was added, and every subsequent 24 hours the media was aspirated and replaced with fresh N2B27. Signalling modulation in gastruloids was performed through addition of small molecule ligands or activators/inhibitors as indicated in the text and figure legends (Table 2).

**Table 2:**
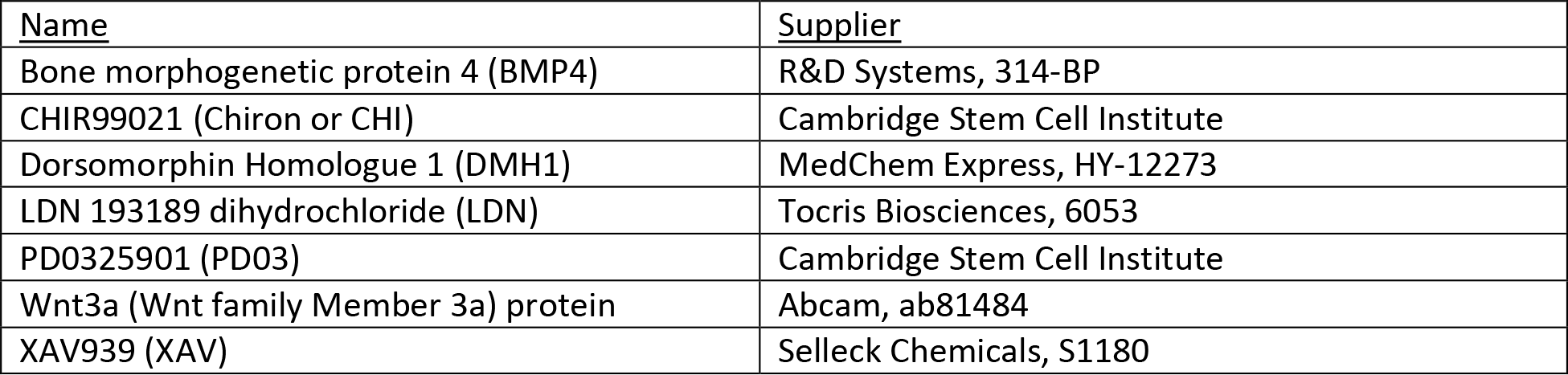
Signalling modulators.

### Immunofluorescence staining

Immunostaining was performed based on a previously published protocol^93^. Gastruloids were collected and washed twice in PBS before fixing in 4% PFA in PBS at 4°C (2h to overnight on an orbital shaker). Three PBS washes to remove the PFA before three washes with blocking buffer PBSFT (10% FBS, 0.2% Triton X-100 in PBS) and blocking in PBSFT for 1-2hr at 4°C on an orbital shaker. Primary antibodies (see Table 3) were added in PBSFT and incubated overnight at 4°C on an orbital shaker. A total of 10 washes with PBSFT were performed before secondary antibodies (diluted 1 in 500) (see Table 4) and Hoechst (Hoechst 33342 trihydrochloride trihydrate, Invitrogen Mol Probes H3570, 10mg/ml solution in water, 16.2mM) at 1 in 800 dilution were added and incubated overnight at 4°C on an orbital shaker. Three PBSFT washes followed by five PBT (0/2% FBS, 0.2% Triton X-100 in PBS) washes were performed before the gastruloids were transferred to ScaleS4 tissue clearing solution (40% D-(-)-sorbitol, 10% Glycerol, 4M Urea, 0.2% Triton X-100, 20% DMSO) in a glass bottom dish and incubated overnight at 4°C on an orbital shaker or mounted on coverslips before imaging.

**Table 3:** Primary antibodies.

| Antibody target | Host species | Supplier | Catalogue number | Dilution |
| --- | --- | --- | --- | --- |
| 5hmC | Rabbit | Abcam | ab214728 | 1 in 100 |
| AP2-gamma | Mouse | Santa Cruz | sc-53162 | 1 in 100 |
| Brachyury | Rabbit | Abcam | ab209665 | 1 in 100 |
| DAZL | Rabbit | Abcam | ab215718 | 1 in 200 |
| E-cadherin | Mouse | Abcam | ab76055 | 1 in 100 |
| E-cadherin | Rat | Takara | M108 | 1 in 100 |
| EpCAM | Rabbit | Abcam | ab221552 | 1 in 100 |
| FoxA2 | Rabbit | Abcam | ab108422 | 1 in 100 |
| FoxC1 | Rabbit | Abcam | ab223850 | 1 in 200 |
| GATA4 | Rabbit | Abcam | ab84593 | 1 in 200 |
| GCNA1 (Tra98) | Rat | Abcam | ab82527 | 1 in 200 |
| GFP | Chicken | Abcam | ab13970 | 1 in 2000 |
| Histone H3K27me3 | Rabbit | Abcam | ab192985 | 1 in 100 |
| Nanog | Rat | ThermoFisher Scientific | 14-5761-80 | 1 in 250 |
| Nanog | Rabbit | Abcam | ab214549 | 1 in 100 |
| N-cadherin | Mouse | BD Biosciences | 610921 | 1 in 200 |
| Oct4 | Rabbit | Abcam | ab200834 | 1 in 200 |
| PECAM1 (CD31) | Rat | BD Biosciences | 557355 | 1 in 200 |
| PhosphoERK | Rabbit | Cell Signalling Technology | 4370 | 1 in 100 |
| PhosphoSMAD1/5/8 | Rabbit | Cell Signalling Technology | 13820 | 1 in 100 |
| Stella (DPPA3) | Goat | R&D Systems | AF2566 | 1 in 50 |
| Sox2 | Rabbit | Abcam | ab92494 | 1 in 200 |
| Sox17 | Rabbit | Abcam | ab224637 | 1 in 100 |
| CDX2 | Rabbit | Abcam | ab76541 | 1 in 200 |

**Table 4:** Secondary antibodies and primary conjugate.

| Antibody target | Antibody species/type | Supplier | Catalogue number | Dye |
| --- | --- | --- | --- | --- |
| Chicken IgY | Goat | Abcam | ab150173 | Alexa 488 |
| Mouse IgG | Donkey | ThermoFisher | A10037 | Alexa 568 |
| Mouse IgG | Goat | ThermoFisher | A21236 | Alexa 647 |
| Rabbit IgG | Donkey | ThermoFisher | A21206 | Alexa 488 |
| Rabbit IgG | Donkey | ThermoFisher | A10042 | Alexa 568 |
| Rabbit IgG | Donkey | ThermoFisher | A31573 | Alexa 647 |
| Rat IgG | Donkey | ThermoFisher | A21208 | Alexa 488 |
| Rat IgG | Donkey | Abcam | ab150153 | Alexa 647 |
| PECAM1(CD31) | Rat | BD Biosciences | 553373 | Phycoerythrin (PE) |

For the 5hmC immunostaining the gastruloids were treated with 1N HCl for 1 hour at room temperature to expose the DNA prior to primary antibody addition. For the phosphorylation antibodies, PBS in the solution buffers was replaced with TBS.

### Imaging

Confocal imaging was performed with either a Zeiss LSM770 or LSM880 Inverted confocal microscopes, using a Plan-Apochromat 20x/0.8 DICII air objective, imaging 6µm Z sections. Data was captured using the Zen software (Carl Zeiss Microscopy Ltd) and images were processed using ImageJ (FIJI)^94^ to generate Z slice section images or Z max projections. Hoechst channel when not shown was used to trace gastruloid outlines to show morphology.

Live imaging was carried out in environmental control units (humidified, 5% C0_2_, 37°C) using either widefield Nikon Inverted Eclipse Ti2 microscopes (15x or 20x ELWD objectives, GFP/YFP/mCherry triple filter) operated by open-source Micro manager software (Vale lab, UCSF, USA) or a Zeiss LSM880 NLO Invert multi-photon microscope (20x objective) operated by Zen software. The Chameleon laser in the multi-photon microscope was tuned to 880nm, with filter 515/30 and 450/80 to detect GFP and CFP and CFP only respectively. Images were captured in single plane every 20 minutes for over 14 hours on the Nikon and Z stacks taken every 30 minutes for over 18 hours on the multi-photon.

### Image analysis

Expression profiles were generated in ImageJ by drawing a segmented line (120 width for whole gastruloid profiles or 20 width for DAZL/NANOG cells) from posterior to anterior of Z max projections of the gastruloids (also used to determine length of gastruloids with ‘measure’ function), plotting the fluorescence profile using the ‘Plot Profile’ function then normalising both the length and signal (against Hoechst) before plotting in Prism (GraphPad) software.

Cell counting (Parameter option: cell size = 8) and gastruloid volume calculations were performed using the IMARIS software (Oxford Instruments), with gastruloid volumes calculated by creating a surface (surface smoothing 1.5, threshold 800-2000) on the Hoechst channel. Cell tracking and co-expression was also performed using IMARIS software. Gastruloid tissue features and PGCLC clusters were assessed by eye in ImageJ. Means, standard deviations and significance (unpaired t-test with Welch’s correction) were calculated in Prism.

### Doubling time calculations

The doubling time of the PGCLCs was calculated based upon the mean cell numbers at each time point, using the following equations to first calculate the growth rate, then the doubling time between time points:

Growth rate (Gr) (%) = ((current cell no. – previous cell no./previous cell no.) x 100
Doubling time (per 24 hours) = (log(2)/log(1 + Gr/100)) x 24

### Flow cytometry and cell sorting

Gastruloids were collected and washed twice in PBS before incubating at room temperature for 8 mins in Trypsin-EDTA before quenching with 10% FBS in PBS. Cells were pelleted at 230xg for 5mins before resuspending in filtered 1% FBS in PBS. Cell solution was passed through tube filter (35µm) then counted and divided into tubes before antibody addition. Incubated at 4°C on rotator for 1 hour then centrifuged at 800 rpm for 5 mins at 4°C. Supernatants were aspirated and sample washed with filtered 1% FBS in PBS before repeating wash and transferring to chilled flow tubes. Cells were applied to a BD FACSAria™ Fusion III (BD Biosciences) performed by the Francis Crick Flow Cytometry Science and Technology Platform (STP) staff. Data analysis was performed using FlowJo (BD Biosciences) software.

Cells were sorted on a BD FACSAria™ Fusion III (BD Biosciences) performed by the Francis Crick Flow Cytometry Science and Technology Platform (STP) staff. Sorting was based on the reporters Stella:eCFP, Blimp1:mVenus or PECAM1-PE and SSEA1-A647 antibodies. Sorted cells were transferred to DNA low-bind tubes and centrifuged at 300 rcf for 5 mins at 4°C. The supernatant was aspirated and, using cut tips, 1ml of chilled PBS pipetted up and down 10 times. This was repeated twice more, and after the final centrifugation step the cells were resuspended in 200µl chilled PBS then 800µl of chilled 100% methanol added dropwise and with stirring. Fixed cells were stored at -80°C until ready for 10x preparation for scRNAseq.

### Single-cell 10x sequencing

The sorted, fixed and frozen cells were thawed on ice for 5 minutes before centrifugation at 1,000 xg for 5 minutes (at 4°C). Supernatant was carefully aspirated without disturbing the pellet before resuspending the pellet in the appropriate volume of Wash-Resuspension buffer; 3x SSC Buffer (Invitrogen, 15557-044) supplemented with 0.04% Bovine Serum Albumin (BSA) (Invitrogen, AM2616), 1mM DL-Dithiothreitol solution (DTT) (Sigma-Aldrich, 646563) and 0.2U/µl Protector RNase inhibitor (Roche, 3335399001) to give 1000 cells/µl in 50µl or minimum volume of 50µl if not possible to obtain that concentration.

Quality control on the cells and counts were performed on a Luna FX7 cell counter (Logos biosystems) prior to applying to 10x Chromium library preparation performed according to manufacturer’s instructions by the Advanced Sequencing Facility staff at the Francis Crick Institute. Single cell libraries of 100 bp paired-end reads were pooled and sequenced using Illumina NovaSeq 6000, carried out by the Advanced Sequencing facility at the Francis Crick Institute.

### scRNA-seq Analysis

FastQ files were quantified into expression matrices using Cell Ranger (6.1.2) using the 10x-provided refdata-gex-mm10-2020-A index. Seurat (4.0.3) objects were created using the filtered matrix for each sorted population in R 4.1.1. Each population was filtered according to the number of reads, features and proportion of mitochondrial expression to remove low-quality cells. Quality-controlled datasets were integrated into a single “120h” dataset with Seurat^95^. Datasets were scaled, projected and clustered using the first 10 principal components for each sorted population or 15 for the integrated dataset.

Published datasets were reprocessed using Seurat 4.0.3 from either the counts matrix of a Seurat object or output of Cell Ranger. The Zhao et al. data was subset to retain only cells whose author-determined cell type included “PGC”. Where possible, the same cell barcodes, variable features and dimensionality were used when reprocessing the published datasets and any published cell metadata was included. Qualitative comparison between the published and recalculated UMAPs reassured us that the structure in the reference data was preserved in our reprocessed objects. For visualisation, UMAP coordinates were reflected to preserve left-to-right time progression, where possible.

Reference and query datasets were subsequently analysed using Seurat^96^ to transfer labels of published data onto the query data and embed the query data into the reference UMAP. We first used the van den Brink dataset as a query and transferred the cell type label onto the 120h dataset which was subsequently filtered for cells that were most-likely PGC-like. The Zhao et al. dataset^66^ was used as a reference for the 120h PGC-like and Ramakrishna et al.^29^ PGC cells identified in the publication as “cluster 5 excluding E10.5”. From this comparison, both cell type labels (“cell type 1” and “cell type 2”) as well as time point were transferred.

## Data Availability

Raw and processed scRNA-seq data for each sorted population are deposited and publicly available in the Gene Expression Omnibus (GEO) at NCBI under accession GSE228406. Processed data includes both filtered and raw expression matrices output by Cell Ranger.

## Acknowledgements

We would like to thank A. Surani, M. Saitou, C. Schroeter and T. Rodriquez for kindly providing cell lines used in this manuscript. We also thank A. Martinez Arias for his support, advice and feedback throughout this project, as well as F. Cermola for feedback on preliminary experiments, C. Mulas and K. Jones for their advice and expertise in CRISPR/Cas9 gene editing and I. Rodriquez Polo for feedback on the manuscript. In addition, the authors gratefully acknowledge the Francis Crick Light Microscopy, Flow Cytometry, Advanced Sequencing and Bioinformatics and Biostatistics Science and Technology Platforms (STPs) for their support and assistance in this work. C.B.C. is an employee of Abcam Ltd. C.B.C., C.B., P.B-B. and N.M. are funded by the Francis Crick Institute which receives its core funding from Cancer Research UK (CC2186), the UK Medical Research Council (CC2186), and the Wellcome Trust (CC2186). P.B-B. and J.N. were funded by a Wellcome Trust Strategic Award (105031/D/14/Z).

