## Supplementary File for "Gastruloid-derived Primordial Germ Cell-like Cells (Gld-PGCLCs) develop dynamically within integrated tissues"

#### Supplementary Figures

**Supplementary Figure 1: Characterisation of Gld-PGCLCs and endodermal tracts. (A)** Z-slice (left) and maximum projection (right) of Blimp1-GFP gastruloid at 120h. **(B-D)** Expression of endodermal markers, FOXA2 **(B)** and SOX17 in Blimp1-GFP **(C)** and BVSC **(D)** gastruloids. **(E-G)** Expression of EpCAM in Blimp1-GFP **(E)** and BVSC **(F)** gastruloids. Cyan arrowheads, STELLA+ cells. **(G)** Z slice showing internal localisation of the EPCAM+, E-CADHERIN (E-Cad)+ tubular structure (left) and z-projection (right). **(H)** Quantification of number of cells co-expressing each pair of proteins, as indicated by colour scale. **(I)** Heterogeneous co-staining of STELLA with OCT4 and AP2γ in Blimp1-GFP gastruloids. White arrowhead, example of a STELLA+ AP2γ- cell. Insets, higher magnification images; Dashed line, morphological gastruloid outline from Hoechst staining; Dotted line, magnification region. Scale bars, 100 µm.

**Supplementary Figure 2: Time course analysis of Gld-PGCLCs as indicated by PECAM1 staining. (A)** Time course of BVSC and Blimp1-GFP gastruloids from 24-144h, showing co-expression of PGCLC markers. Cyan arrowheads, STELLA+ cells; Yellow arrowheads, AP2γ+ cells; Green arrowheads, STELLA+ AP2γ+ cells. Insets, higher magnification images; Dashed line, morphological gastruloid outline from Hoechst staining; Dotted line, magnification region. Scale bars, 100 µm. **(B)** Quantification of BVSC cells expressing protein indicated that also express PECAM1, as percentages, at 120h. **(C)** Flow cytometry analysis of Stella:eCFP from BVSC line and PECAM1, where dot-plot colour-bar shows data density. Numbers in graph show percentage occupancy of each quadrant.

**Supplementary Figure 3: Expression of cell membrane markers on Gld-PGCLCs and endodermal tract cells. (A)** Expression of E-Cadherin (E-Cad) in BVSC and Blimp1-GFP gastruloids. Red arrowheads, E-CAD on Gld-PGCLCs and neighbouring cells. **(B)** Expression of EpCAM in BVSC gastruloids. Red arrowheads, expression of EpCAM on Gld-PGCLCs and neighbouring cells. Insets, higher magnification images; Dashed line, morphological gastruloid outline from Hoechst staining; Dotted line, magnification region. Scale bars, 100 µm.

**Supplementary Figure 4: Anterior Gld-PGCLC cluster associated gene expression. (A-C)** Expression of GATA4 in anterior regions adjacent to Gld-PGCLCs, in Blimp1-GFP **(A, B)** and BVSC **(C)** gastruloids. **(D)** A published gastruloid dataset shows expression of *Gata4* and *Cxcl12* genes (left), particularly localised to anterior end as evidenced by tomo-sequencing (right). Individual lines represent different sample replicates, black line represents the average profile, and grey ribbon represents the standard deviation (Stdev). See [1] for further details. **(E)** Expression of GCNA1 particularly in AP2γ+ Gld-PGCLCs. Insets, higher magnification images; Dashed line, morphological gastruloid outline from Hoechst staining; Dotted line, magnification region. Scale bars, 100 µm.

**Supplementary Figure 5: Analysis of single-cell transcriptomics. (A)** UMAP of Gld-PGCLCs from all 3 sorted populations, showing 8 distinct clusters according to set parameters (see Methods). **(B)** Gene expression within clusters, grouped by biological categories. Shading indicates the potential identity of clusters: cluster 0-4 as PGC-like, cluster 5 as Meso+Somiticlike, cluster 6 as Endoderm-like and cluster 7 as endothelium-like. **(C)** Module Score information for selected gene signatures across the UMAP representation. **(D)** Cell label transfer from published gastruloid dataset[1], showing general concordance with gene expression annotation. **(E)** Quantification of cells assigned to identities by label transfer with the van den Brink dataset.

**Supplementary Figure 6: BMP signalling and Gld-PGCLCs. (A)** Effect of BMP4 addition on Blimp1-GFP gastruloids at time indicated. **(B)** Quantification of AP2γ+ cell counts in Blimp1-GFP gastruloids at 120h. **(C-D)** Localisation of phospho-SMAD1/5/8 in BVSC **(C)** and Blimp1-GFP **(D)** gastruloids. Yellow arrowheads, Gld-PGCLCs. Insets, higher magnification images; Dashed line, morphological gastruloid outline from Hoechst staining; Dotted line, magnification region. Scale bars, 100 µm.

**Supplementary Figure 7: Effect of reduced BMP signalling on Gld-PGCLCs. (A)** Inhibition of BMP signalling by DMH1 or LDN1 at times indicated. **(B)** Effect of BMP signalling modulation in the absence of Wnt signalling through Chi exposure. **(C)** Quantification of AP2γ+ cell counts in Blimp1-GFP gastruloids at 120h following DMH1 exposure. **(D)** Quantification of AP2γ+ cell counts in BVSC and Blimp1-GFP gastruloids (Wildtype) or BMPR1A-/- at 120h. **(E-I)** BMPR1A-/- gastruloids show presence of markers of mesoderm, FOXC1, **(E),** neural, SOX2, **(F),** endoderm, FOXA2 and SOX17**, (G, H)** and ectoderm, N-Cadherin **(I).** Red arrowheads, SOX2+ NANOG- ectodermal cell examples. White arrowheads, SOX2+ N-CAD- Gld-PGCLC examples. Insets, higher magnification images; Dashed line, morphological gastruloid outline from Hoechst staining; Dotted line, magnification region. Scale bars, 100 µm.

**Supplementary Figure 8: Wnt signalling and Gld-PGCLCs. (A-B)** Maximum projection of gastruloids following shifts in Chi exposure, at timepoint and concentrations indicated. **(C)** Quantification of AP2γ+ cell counts in BVSC gastruloids at 120h following Chi exposure at timepoints indicated. **(D)** Maximum projection of BVSC gastruloids following increase in Chi concentration at timepoints indicated. **(E)** Quantification of AP2γ+ cell counts in BVSC gastruloids at 120h following Chi concentration modulation at 48-72h. **(F)** Maximum projection of Blimp1-GFP gastruloids following increase in Chi concentration at timepoints indicated. **(G)** Quantification of AP2γ+ cell counts in Blimp1-GFP gastruloids at 120h following Chi concentration modulation at 48-72h. **(H)** Maximum projection of BVSC gastruloids following Wnt signalling modulation by ligand application, Wnt3a, or inhibitor exposure, XAV, in the absence of Chi pulse. **(I)** Quantification of AP2γ+ cell counts in BVSC gastruloids at 120h following Wnt signalling modulation, at timepoint and concentrations indicated. Dashed line, morphological gastruloid outline from Hoechst staining; Dotted line, magnification region. Scale bars, 100 µm.

**Supplementary Figure 9: FGF signalling and Gld-PGCLCs. (A)** Z slice of phospho-ERK in BVSC (left) and Blimp1-GFP (right) gastruloids at onset of Gld-PGCLC specification. Yellow arrowheads, AP2γ+ pERK- cells. **(B)** Maximum projection of BVSC gastruloids with inhibition of FGF signalling by exposure to PD03 at time and concentration indicated, in the absence of Chi pulse. **(C)** Quantification of AP2γ+ cell counts in BVSC gastruloids at 120h following PD03 exposure at 24-48h, in the absence of Chi pulse. **(D)** Maximum projection of BVSC gastruloids following FGF signalling modulation by PD03 at timepoints indicated. Insets, higher magnification images; Dashed line, morphological gastruloid outline from Hoechst staining; Dotted line, magnification region. Scale bars, 100 µm.

### Supplementary Movies

**Supplementary Movie 1:** Widefield imaging of two BVSC gastruloids from 98h after aggregation, imaged at 20 minute intervals. Cyan, Blimp1-GFP; Yellow, Stella:eCFP. Scale bar = 100 um.

**Supplementary Movie 2:** Multiphoton live-imaging movie of a BVSC gastruloid from 118h after aggregation, imaged at 30 minute intervals. Cyan, Blimp1-GFP; Yellow, Stella:eCFP. Scale bar = 100 um.

### Supplementary Tables

**Supplementary Table 1:** Table of gastruloid lengths and volumes, from BVSC and Blimp1-GFP gastruloids at 120h and 144h.

120h

| | Gastruloid type | |
|---|---|---|
| Gastruloid measurements | BVSC | Blimp1-GFP |
| Mean Length (μm) (± SD) | 607.00 ± 122.20 | 933.07 ± 106.00 |
| Mean volume ($\mu m^3$) (± SD) | $0.66 \times 10^7 \pm 2.62 \times 10^6$ | $2.17 \times 10^7 \pm 5.90 \times 10^6$ |
| Number of samples | 32 | 16 |

144h

| | Gastruloid type | |
|---|---|---|
| Gastruloid measurements | BVSC | Blimp1-GFP |
| Mean Length (μm) (± SD) | 301.9 ± 45.53 | 1014 ± 205.5 |
| Mean volume ($\mu m^3$) (± SD) | $0.33 \times 10^7 \pm 0.24 \times 10^6$ | $1.86 \times 10^7 \pm 5.82 \times 10^6$ |
| Number of samples | 24 | 18 |

**Supplementary Table 2:** Table of number of cells per Gld-PGCLC cluster in 144h gastruloids. Gast, Gastruloid replicate.

| | Gast 1 | | | | | Gast 2 | | | | | Gast 3 | Gast 4 | | | Gast 5 | | | Gast 6 | Gast 7 | | | | |
|---|---|---|---|---|---|---|---|---|---|---|---|---|---|---|---|---|---|---|---|---|---|---|---|
| Cluster no. | 1 | 2 | 3 | 4 | 5 | 6 | 7 | 8 | 9 | 10 | 11 | 12 | 13 | 14 | 15 | 16 | 17 | 18 | 19 | 20 | 21 | 22 | 23 |
| AP2γ + PGC marker | 0 | 2 | 1 | 4 | 3 | 6 | 4 | 7 | 6 | 0 | 0 | 4 | 3 | 8 | 4 | 2 | 5 | 7 | 5 | 6 | 5 | 6 | 8 |
| AP2γ + Blimp1 | 5 | 6 | 6 | 1 | 3 | 0 | 0 | 5 | 0 | 0 | 12 | 0 | 0 | 0 | 0 | 0 | 0 | 0 | 0 | 0 | 0 | 0 | 0 |
| PGC marker + Blimp1 | 0 | 0 | 0 | 0 | 0 | 0 | 0 | 0 | 0 | 0 | 0 | 0 | 0 | 0 | 0 | 0 | 0 | 0 | 0 | 0 | 2 | 0 | 0 |
| AP2γ + PGC marker + Blimp1 | 0 | 6 | 3 | 0 | 0 | 3 | 0 | 0 | 8 | 0 | 1 | 4 | 13 | 5 | 6 | 11 | 0 | 2 | 2 | 0 | 3 | 0 | 4 |
| Total | 5 | 14 | 10 | 5 | 6 | 9 | 4 | 12 | 14 | 0 | 13 | 8 | 16 | 13 | 10 | 13 | 5 | 9 | 7 | 6 | 10 | 6 | 12 |
| Total cells in cluster | 9 | 14 | 11 | 7 | 10 | 9 | 6 | 15 | 14 | 10 | 13 | 8 | 16 | 13 | 10 | 13 | 5 | 9 | 13 | 6 | 10 | 7 | 12 |
| PGC marker = DAZL/PECAM1 | | | | | | | | | | | | | | | | | | | | | | | |

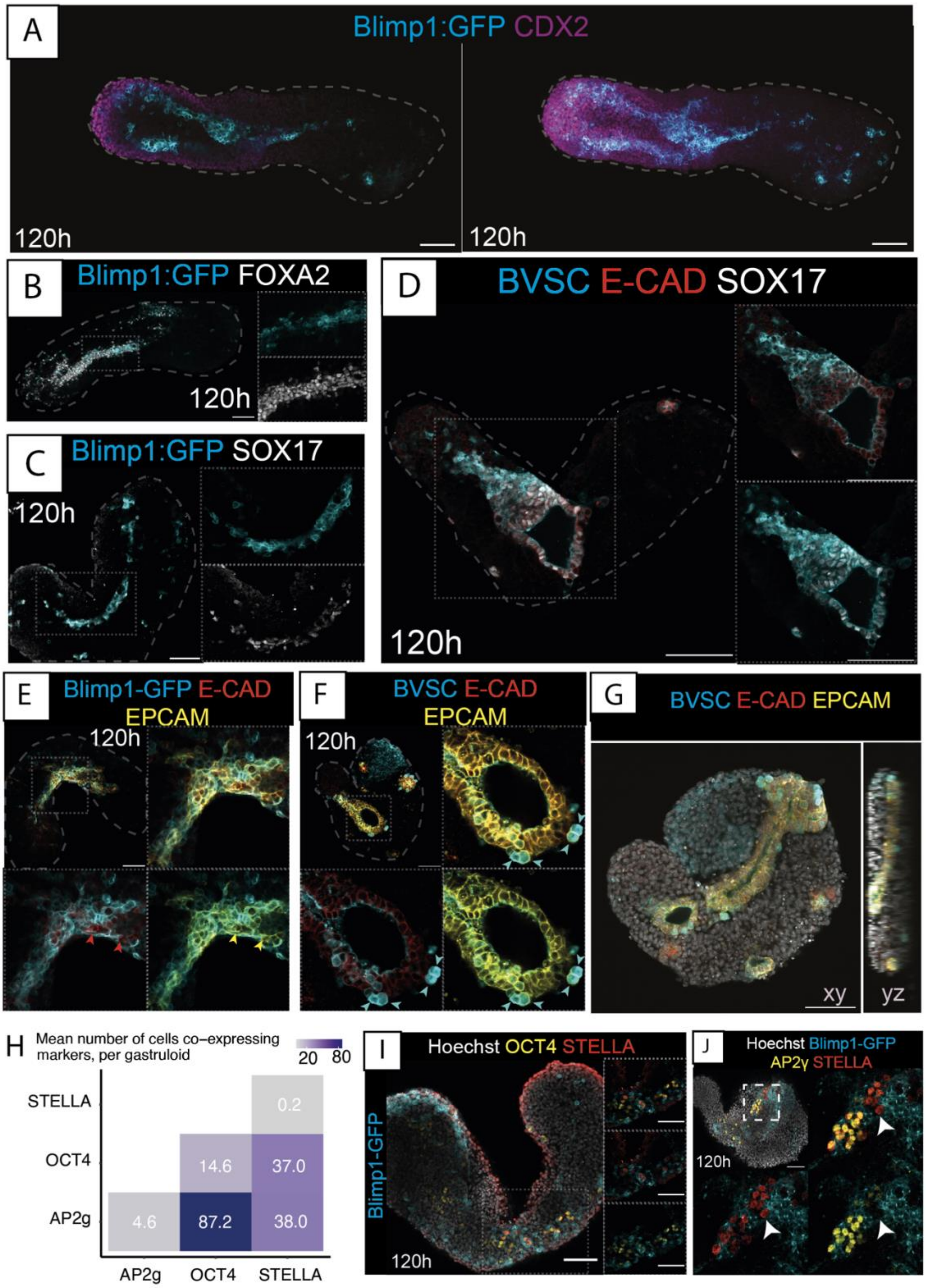

Supplementary Figure 1

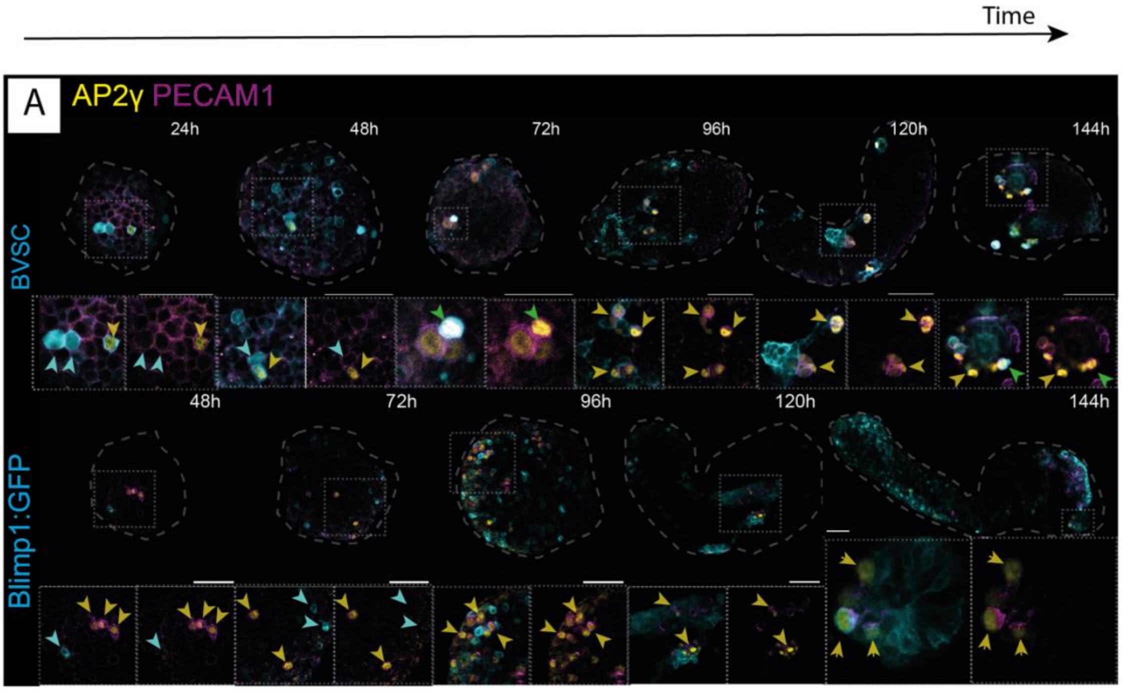

B

| | Percentage co-expression with PECAM1 |
|---|---|
| **AP2γ** | 96.65 |
| **OCT4** | 95.83 |
| **STELLA** | 98.30 |

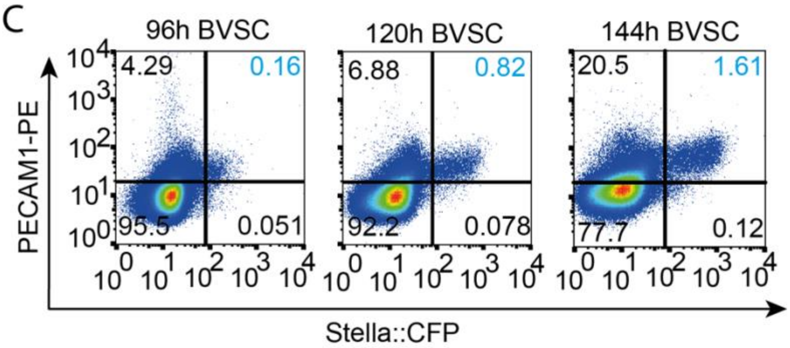

Supplementary Figure 2

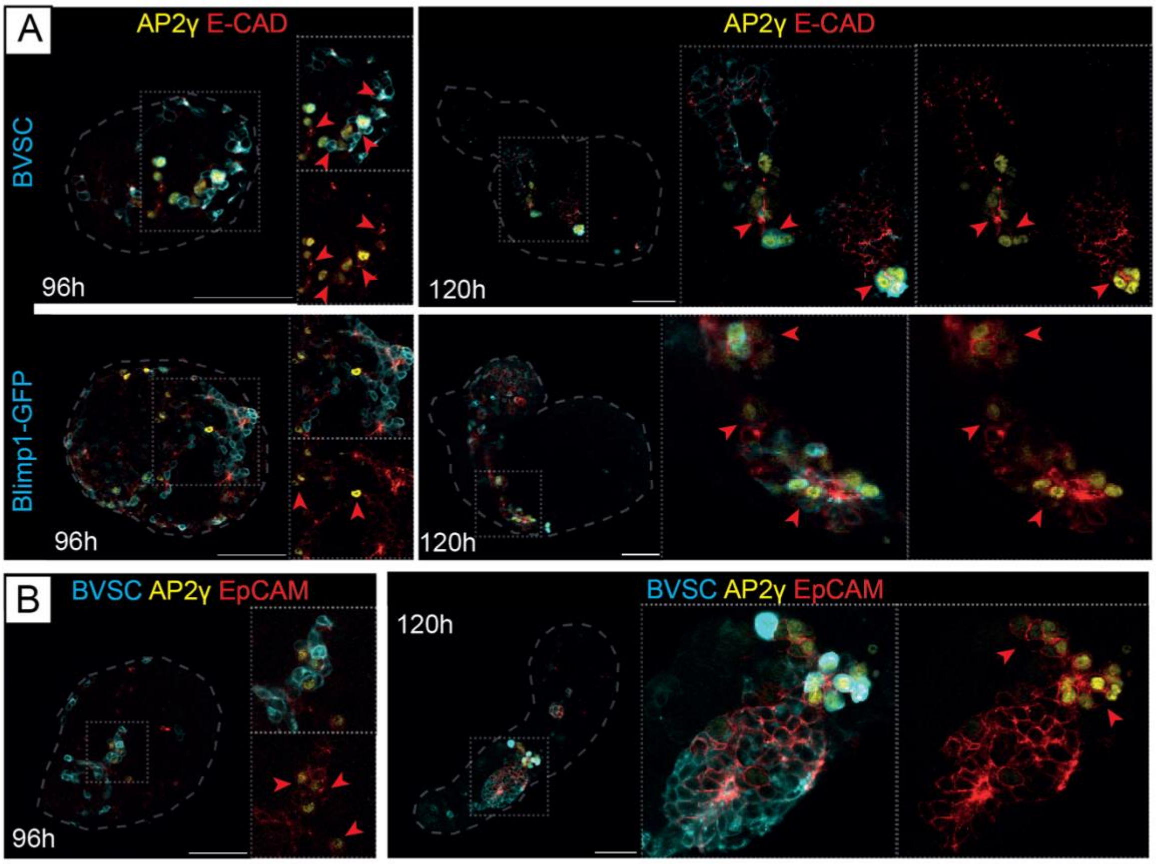

Supplementary Figure 3

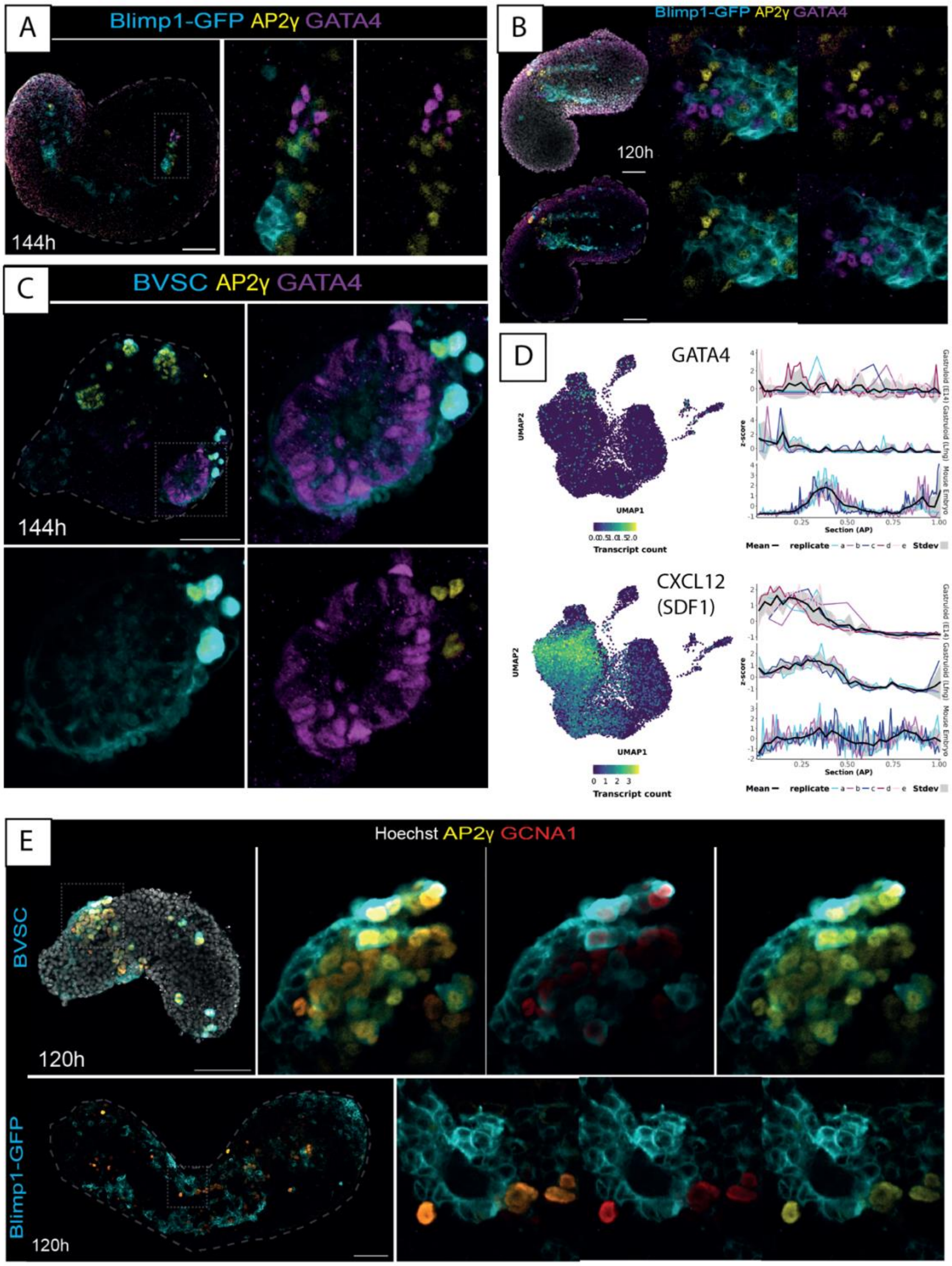

Supplementary Figure 4

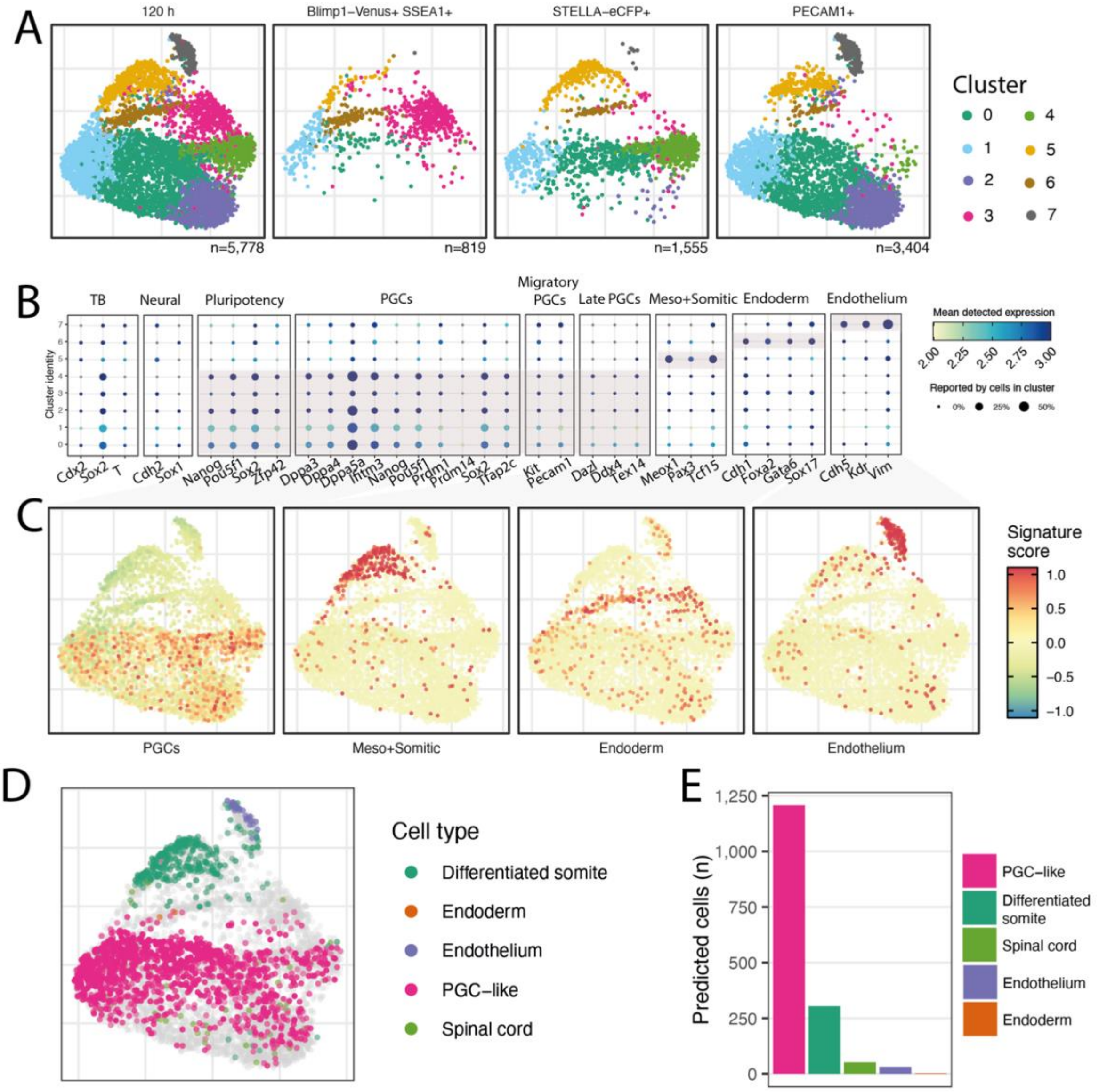

Supplementary Figure 5

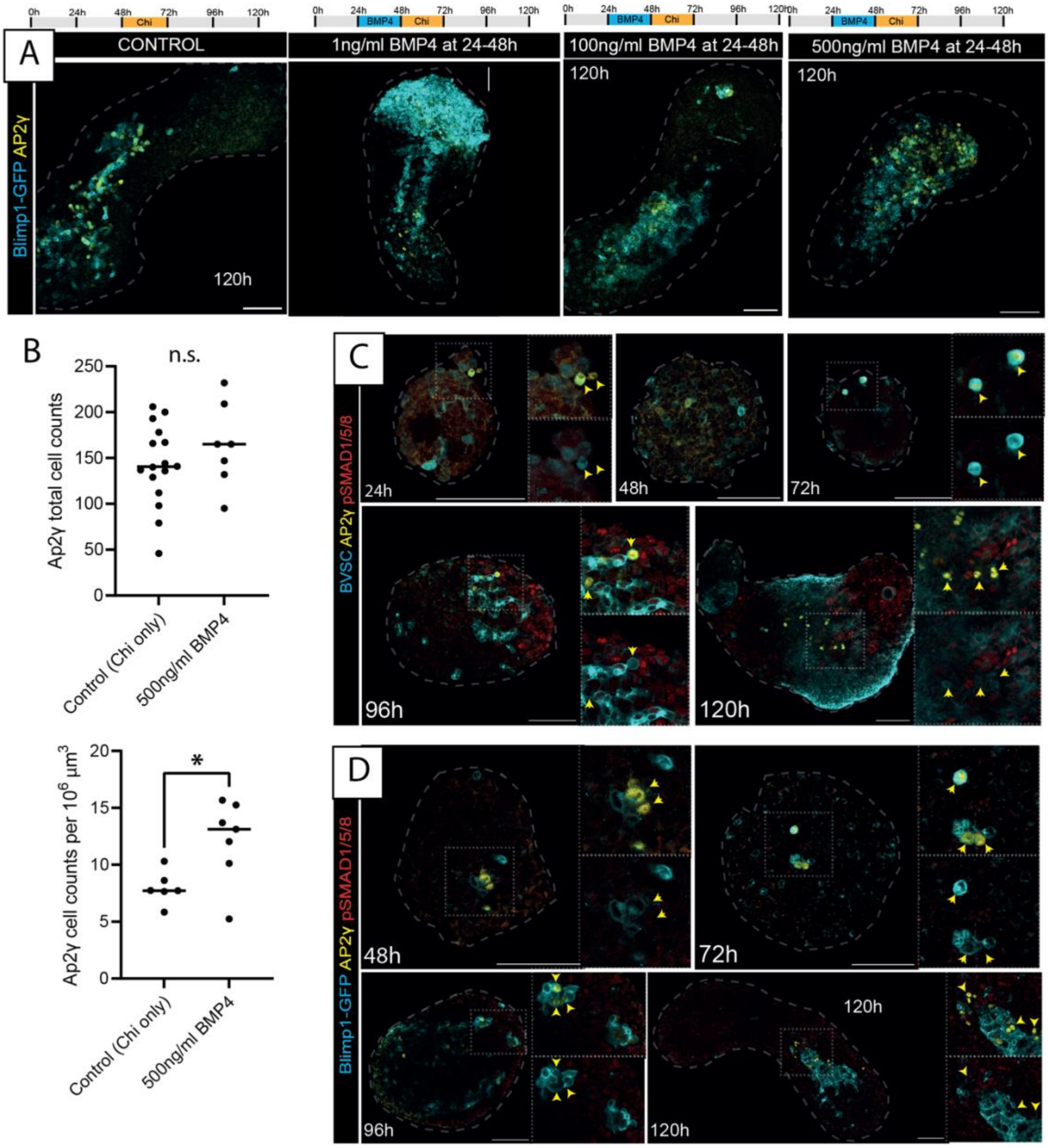

Supplementary Figure 6

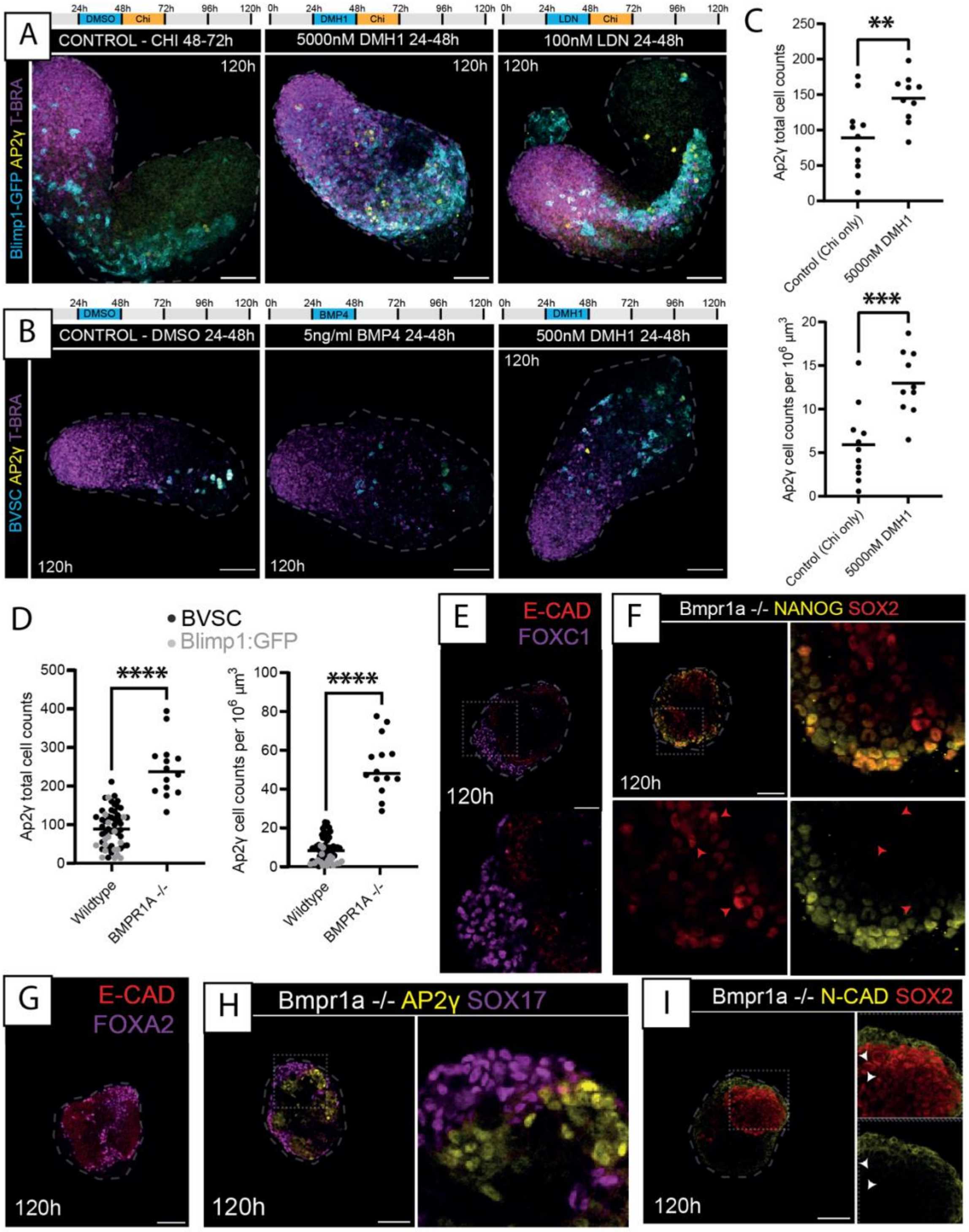

Supplementary Figure 7

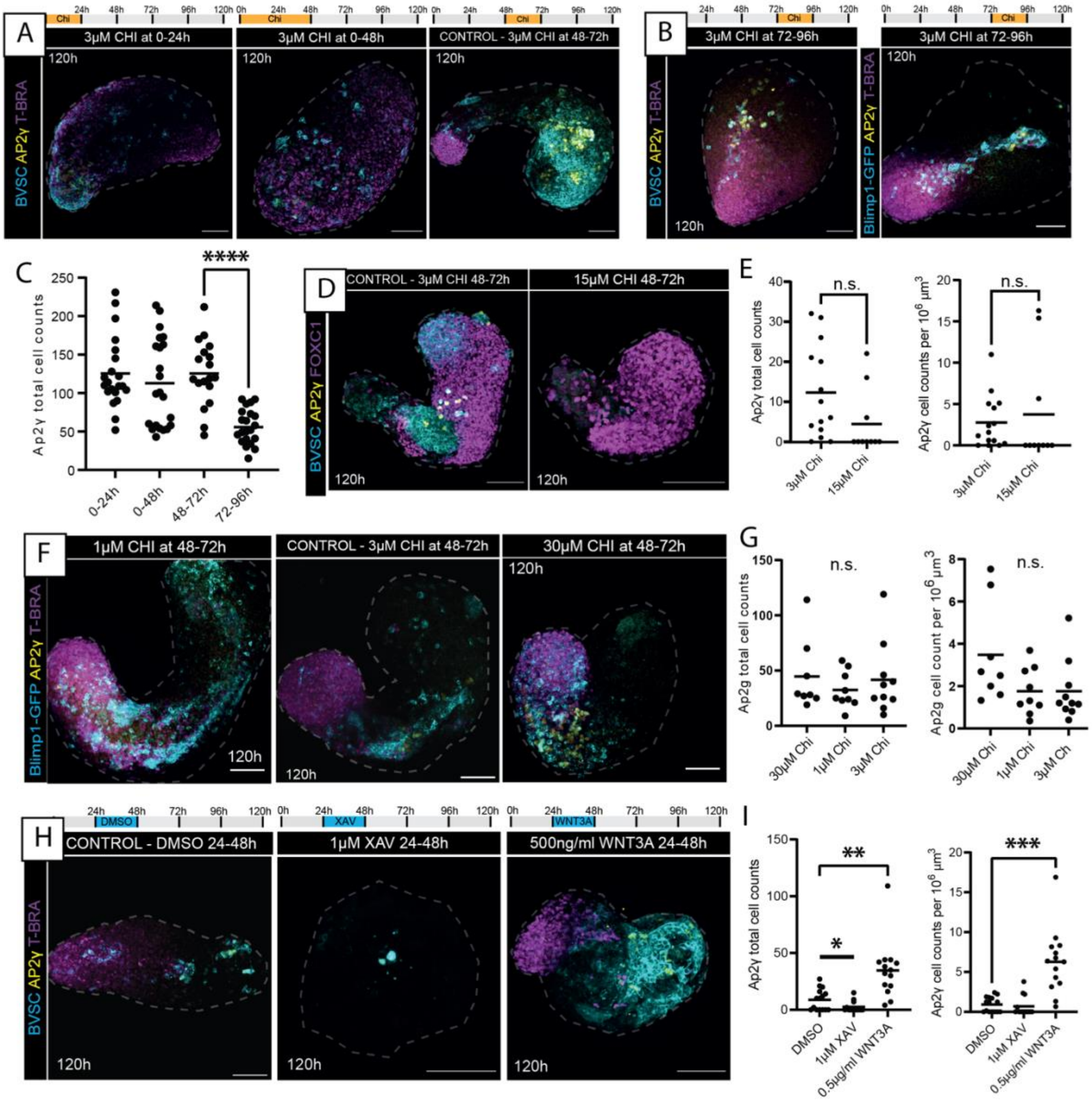

Supplementary Figure 8

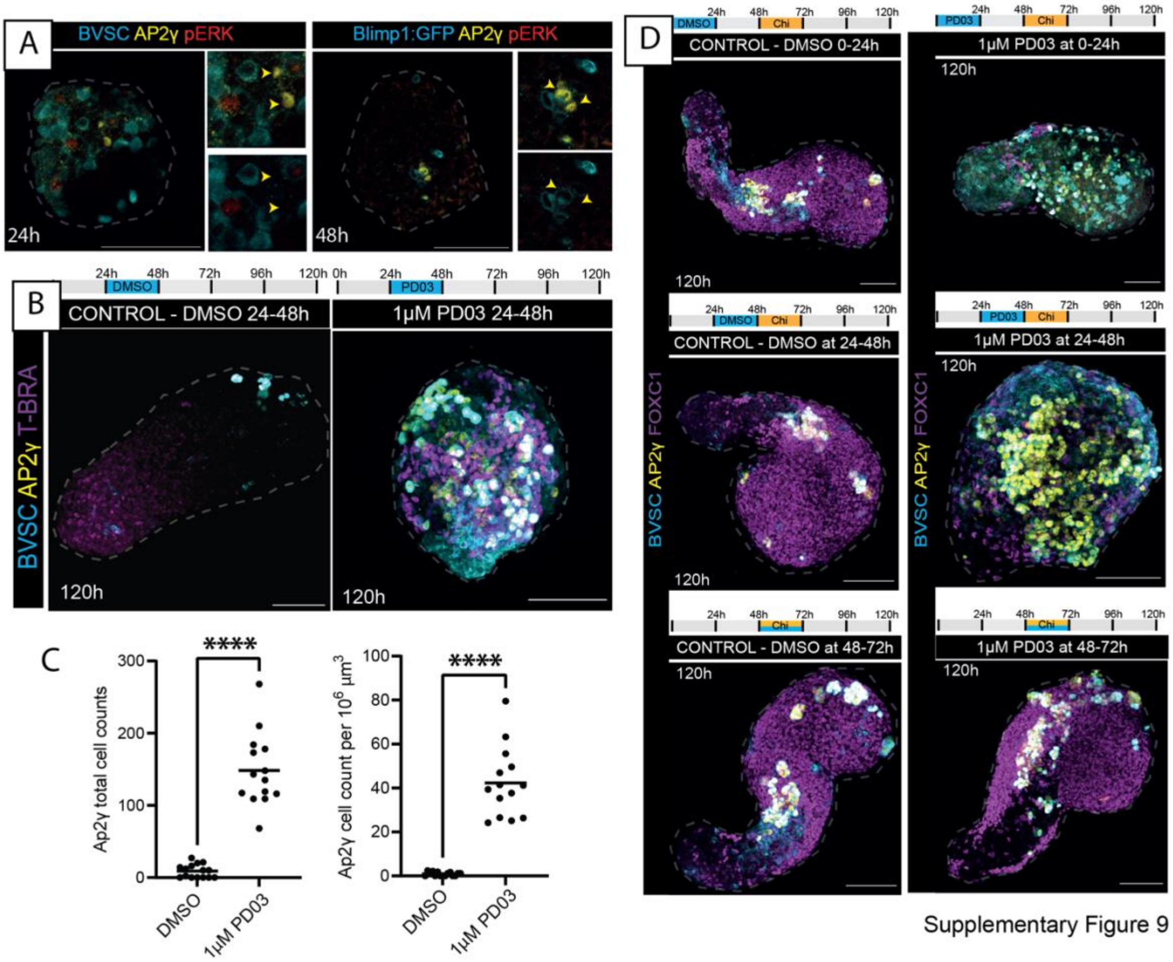

Supplementary Figure 9

**1**. van den Brink, S. C. *et al.* (2020) Single-cell and spatial transcriptomics reveal somitogenesis in gastruloids. *Nature* 582, 405-9 https://doi.org/10.1038/s41586-020-2024-3.
